# BRN2 is a non-canonical melanoma tumor-suppressor

**DOI:** 10.1101/2021.01.13.426554

**Authors:** Michael Hamm, Pierre Sohier, Valérie Petit, Jérémy H Raymond, Véronique Delmas, Madeleine Le Coz, Franck Gesbert, Colin Kenny, Zackie Aktary, Marie Pouteaux, Florian Rambow, Alain Sarasin, Alfonso Bellacosa, Luis Sanchez-del-Campo, Laura Mosteo, Martin Lauss, Dies Meijer, Eirikur Steingrimsson, Göran B Jönsson, Robert A Cornell, Irwin Davidson, Colin R Goding, Lionel Larue

## Abstract

While the major drivers of melanoma initiation, including activation of NRAS/BRAF and loss of *PTEN* or *CDKN2A*, have been identified, the role of key transcription factors that impose altered transcriptional states in response to deregulated signaling is not well understood. The POU domain transcription factor BRN2 is a key regulator of melanoma invasion, yet its role in melanoma initiation remains unknown. Here, we show that *BRN2* haplo-insufficiency is sufficient to promote melanoma initiation and metastasis, acting as a non-canonical tumor suppressor. Mechanistically, BRN2 directly modulates *PTEN* expression, and PI3K signaling, to drive tumor initiation and progression. Collectively our results reveal that somatic deletion of one *BRN2* allele elicits melanoma initiation and progression.

**SIGNIFICANCE:** Here, we report frequent mono-allelic loss of the transcription factor BRN2 in human cutaneous melanoma metastases. We developed a mouse model for Brn2-deficient melanoma based on the most common alterations (*Braf*^*V600E*^ and *Pten* loss) in human melanoma and established the role of Brn2 as a functional regulator of tumor initiation, tumor growth, and the formation of metastases *in vivo*. Mechanistically, BRN2 loss increases PI3K-signaling through PTEN repression, either via MITF induction or not. Overall, we describe a novel tumor suppressor of high prevalence in human melanoma that regulates several steps of *in vivo* melanomagenesis through two previously unknown molecular mechanisms.

## INTRODUCTION

Cancer initiation is triggered by the activation of oncogenic signaling combined with senescence bypass. Yet while many of the typical oncogenes and tumor suppressors that affect cancer initiation have been identified, cancer initiation is likely to be modulated by additional genetic events. Understanding how non-classical driver mutations may impact cancer initiation is a key issue that has been relatively underexplored. Melanoma, a highly aggressive skin cancer, arises through the acquisition of well-defined genetic and epigenetic modifications in oncogenes and tumor suppressors and represents an excellent model system to address this key question.

As a highly genetically unstable cancer type, the initiation of melanoma requires the induction of melanocyte proliferation, which is mediated by several major founder mutations, the most common of which are BRAF^V600E^ and NRAS^Q61K/R 1,2^. However, activation of BRAF or NRAS is insufficient to promote melanoma initiation without senescence bypass mediated by additional founder mutations or expression changes of several genes including p16^INK4A^, β-catenin, PTEN, or MDM4 ^3-7^.

The transcription factor BRN2, also known as POU3F2 and N-OCT3, plays a critical role in neurogenesis and is expressed in a range of cancer types with neural or neuroendocrine origins, in which it drives proliferation, including glioblastoma, neuroblastoma, small cell lung cancer, and neuroendocrine prostate cancer ^8-10^. In the melanocyte lineage, BRN2 is not detected in melanoblasts *in vivo* but is heterogeneously expressed in melanoma ^11-13^. *In vitro* studies have shown that BRN2 expression is induced by a range of melanoma-associated signaling pathways including activation of the mitogen-activated protein kinase (MAPK) pathway downstream from BRAF, the PI3K pathway, the LEF-β-catenin axis, as well as FGF, TNF-α, EDN3 and SCF signaling ^13-16^. Consistent with BRN2 being expressed in a predominantly mutually exclusive pattern with the Microphthalmia-associated transcription factor (MITF) that plays a crucial role in melanoma proliferation, BRN2 is repressed by MITF via miR-211 ^17^. The activation of BRN2 expression in a specific subset of melanoma cells in response to all three major signaling pathways linked to melanoma progression suggests it is likely to have a critical role in disease progression ^18^. Most notably, BRN2 has been associated with slow-cycling, MITF-low cells and identified as a key regulator of melanoma invasion *in vitro* ^12,19^ and in *in vivo* (xenograft) experiments ^20-22^. Mechanistically, the ability of BRN2 to promote invasion has been linked to its ability to control expression of PDE5A-mediated cell contractility, phosphorylation of myosin light chain 2, repression of MITF and PAX3, and cooperation with bi-allelic loss of CDKN2A ^12,19,20,22^.

However, despite abundant information linking BRN2 to melanoma invasiveness *in vitro* and in *in vivo* xenograft experiments, the impact of BRN2 on melanoma initiation and progression *in vivo* has never been assessed.

## RESULTS

### BRN2 loss or low expression correlates with reduced survival and worse prognosis

Although BRN2 has been implicated in melanoma invasiveness, whether and how it may contribute to melanoma initiation or incidence is not understood. To evaluate the prevalence of *BRN2* loss in human <u>sk</u>in <u>c</u>utaneous <u>m</u>elanoma (SKCM), we retrieved copy-number alteration (CNA) data for *BRN2* in SKCM metastases (stage IV) from The Cancer Genome Atlas (TCGA, https://cancergenome.nih.gov/) and used the definition of the GISTIC (Genomic Identification of Significant Targets in Cancer) values of “-1” for mono-allelic loss and “-2” for bi-allelic loss. The *BRN2* locus showed mono-allelic loss in 53% and bi-allelic loss in 2.7% of all patient samples (n = 367, Figure 1A). Only a minority (n = 29 of 367, corresponding to 7.9%) of SKCM samples showed a copy-number gain/amplification for *BRN2* (GISTIC > “+1” and “+2”), which were not further analyzed (data not shown). We screened a panel of human melanoma cell lines available in our laboratory (n = 23) for deletions that affect the *BRN2* locus by comparative genomic hybridization. The *BRN2* locus showed mono-allelic loss in 48% (11 out 23) of the human melanoma cell lines and no bi-allelic loss, comparable to the TCGA-data (Figure S1A, Table S1), and BRN2 mRNA levels were significantly lower in SKCM metastases with bi-allelic BRN2 loss (Figure S2A). The mono- and bi-allelic loss of *BRN2* was frequently associated with a large segmental deletion of the long arm of chromosome 6 (Chr.6q) in SKCM metastases and in our cell-line panel (Figure 1B, Table S2, Figure S1B). Patients carrying the monoallelic loss of *BRN2* in metastases displayed significantly shorter overall survival than those with diploid status (Figure 1C). These results were confirmed using an independent cohort of 118 regional metastases previously described ^23^ (Figure 1D).

**Figure 1.**
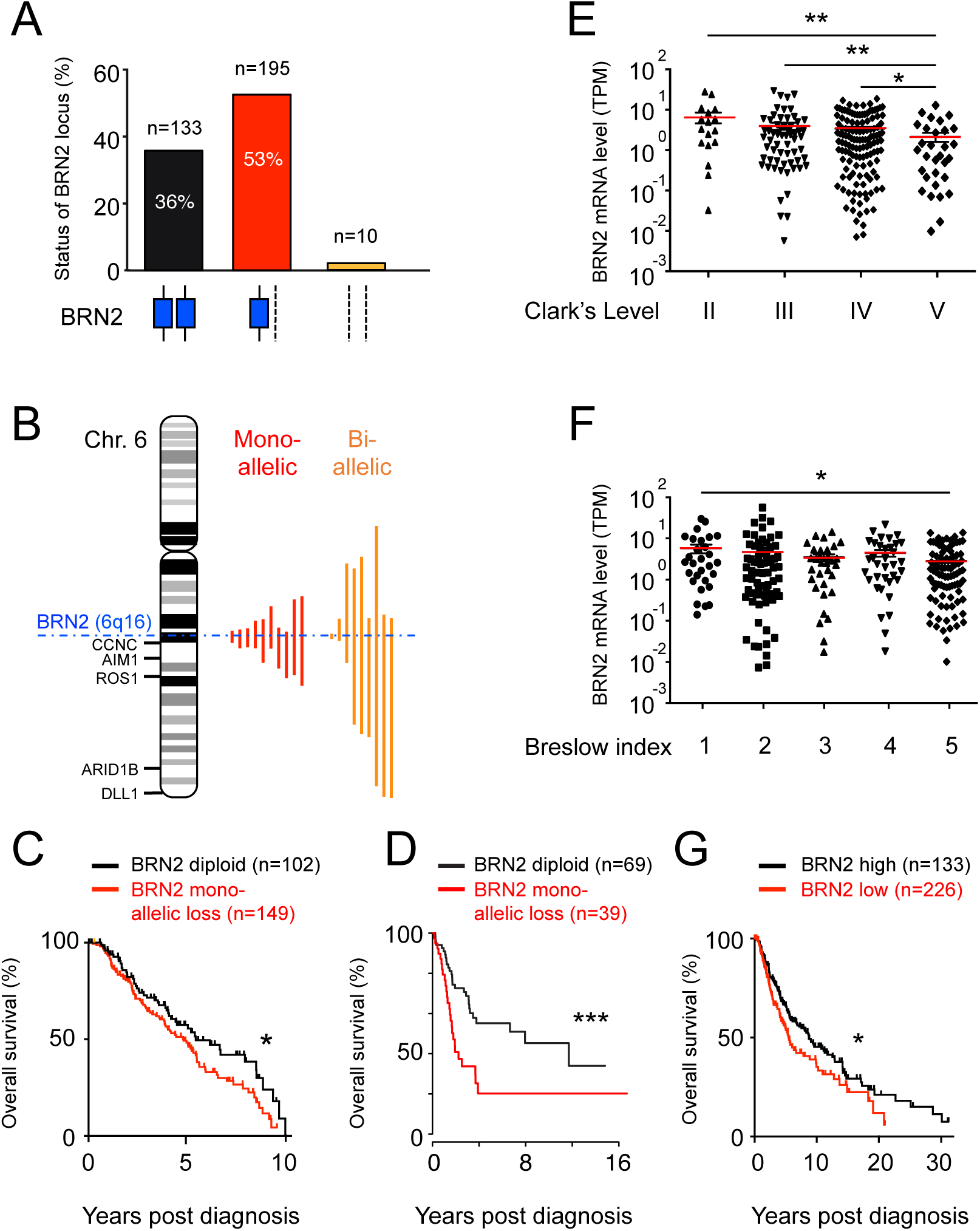
The BRN2 locus is frequently deleted in human melanoma and reduced BRN2 mRNA levels correlate with reduced overall survival and worsening of prognostic factors. (A) Bar graph showing the status of the BRN2 locus in human skin cutaneous melanoma (SKCM) metastases (stage IV). Copy-number alterations (CNAs) were estimated using the GISTIC algorithm. (B) Pictogram showing the extent of segmental deletions that affect the BRN2 locus on Chr.6q16 (dashed blue line) in SKCM metastases. (C) Kaplan-Meier curves comparing 10-year overall survival of SKCM patients diploid for BRN2 (black line, n = 102) or those with mono-allelic (red n = 149). The TCGA CNA-data set was analyzed (n = 309). Diploid vs. Mono-allelic loss: log-rank (Mantel-cox) test, Diploid vs. Bi-allelic loss: Gehan-Breslow-Wilcoxon test (p = 0.027). (D) Kaplan Meier curves comparing SKCM patients with diploid status or mono-allelic loss of BRN2 in 118 regional metastatic melanoma patients (p = 0.003, log-rank test). (E) Correlation between BRN2 mRNA levels and Clark levels of SKCM samples. Significance was determined using the Mann-Whitney test. (F) Correlation between BRN2 mRNA levels and Breslow index in SKCM samples. The means are shown. Significance was determined using the Mann-Whitney test. (G) Kaplan-Meier curves comparing 30-year overall survival of SKCM patients to BRN2 mRNA levels. Log-rank (Mantel-Cox) test (p = 0.03). All data were retrieved from TCGA on May 28, 2017. Significance was defined as * (p < 0.05), ** (p < 0.01), and *** (p≤0.001).

We next assessed the correlation between BRN2 mRNA levels in SKCM metastases and the Clark’s level, Breslow index of matched primary tumors, and melanoma stage, defined by the AJCC ^24,25^. Reduced BRN2 mRNA levels correlated with higher Clark’s level, Breslow index, and AJCC score (Figure 1E, F and Figure S2B). Finally, we assessed the correlation between BRN2 mRNA levels and overall patient survival to evaluate the effect of BRN2 loss on melanoma progression. We established “BRN2-high” and “BRN2-low” patient groups based on RNA-seq data available from the TCGA (BRN2 subgroups defined as BRN2 low (≤ 1 transcript per million reads [TPM]) and BRN2 expressed/high (> 1 TPM), total patients n = 359). Patients in the “BRN2 low” group displayed significantly shorter overall survival than those of the “BRN2 high” group (Figure 1G). Overall, the *BRN2* locus, frequently associated with a large segmental deletion, is lost (mono- and bi-allelic) in ≈60% of human SKCM metastases and correlates with significantly reduced overall survival. Lower levels of *BRN2* mRNA from the bulk tumor correlate with higher Clark’s levels and Breslow indices and reduced overall survival.

### Co-occurrence of BRN2 loss with mono-allelic loss of PTEN

We next determined whether BRN2 loss co-occurs with melanoma driver mutations by examining the TCGA CNA-data set (n = 367) and human melanoma cell-line panel (n = 23). There was no significant correlation between *BRAF* or *NRAS* mutation and *BRN2* loss (mono-or bi-allelic), neither in human melanoma samples nor the human cell-line panel (Figure S3A, B). We then searched for co-occurring CNAs of other known melanoma-associated genes and found that mono-allelic loss of BRN2 co-occurred with mono-allelic loss of PTEN in approximately 40% of the human melanoma samples in TCGA and in the human cell-line panel (Figure S3C, D). We next evaluated the concomitant *BRN2* locus alterations and *BRAF*/*NRAS* mutations and *CDKN2A*/*PTEN* alterations and found the most frequent genetic constellation that co-occurs with *BRN2* loss in melanoma to be *BRAF*^*V600X*^ mutation and mono-allelic *PTEN* loss (Figure S3E).

### Loss of BRN2 drives melanomagenesis *in vivo*

These data suggesting that monoallelic loss of BRN2 might be of importance in melanoma prompted us to evaluate the role of BRN2 in melanomagenesis *in vivo* by examining whether the loss of *Brn2* affects melanoma initiation and/or progression in a mouse model. To this end, we developed an inducible genetically engineered mouse model system for Brn2-deficient melanoma driven by the most common alterations in human SKCM (*Braf*^*V600E*^ and *Pten* loss). Specifically, we used *Tyr::Cre*^*ERt2/°*^; *Braf*^*V600E/+*^ (called Braf from hereon) and *Tyr::Cre*^*ERt2/°*^; *Braf*^*V600E/+*^; *Pten*^*F/+*^ (called Braf-Pten from hereon) mice carrying a tamoxifen-inducible Cre-recombinase under the control of the tyrosinase promoter ^26-28^. This model system allows melanocyte lineage-specific induction of a *BRAF*^*V600E*^ mutation and mono-allelic deletion of *Pten* for Braf-Pten mice. Cre-mediated defloxing leads to activation of the Braf^V600E^ oncogene, inducing nevus and spontaneous melanoma formation in Braf mice, reproducing many of the cardinal histological and molecular features of human melanoma ^29^. Bi-allelic and mono-allelic loss of *PTEN* reduces tumor latency in Braf^V600E^- and NRAS^Q61K^-driven mouse melanoma models ^3,30^.

Using these models, we studied the effect of Brn2 haplo-insufficiency on *in vivo* melanomagenesis by introducing the floxed Brn2 locus into the genome by appropriate crossings (Figure S4A) ^31^. Specifically, we generated the mouse lines *Tyr::Cre*^*ERt2/°*^; *Braf*^*V600E/+*^; *Brn2*^*+/+*^ (Braf-WT), *Tyr::Cre*^*ERt2/°*^; *Braf*^*V600E/+*^; *Brn2*^*F/+*^ (Braf-Brn2), *Tyr::Cre*^*ERt2/°*^; *Braf*^*V600E/+*^; *Pten*^*F/+*^; *Brn2*^*+/+*^ (Braf-Pten-WT) and *Tyr::Cre*^*ERt2/°*^; *Braf*^*V600E/+*^; *Pten*^*F/+*^; *Brn2*^*F/+*^ (Braf-Pten-Brn2). Cre-mediated defloxing of Braf, Pten, and Brn2 loci was induced by topical application of tamoxifen during the first three days after birth (Figure S4B). All mice were monitored for the number of tumors/mice, and the growth rate of the first tumor.

Braf-Brn2 mice showed no differences in the number of tumors/mouse or the tumor growth rate from those of Braf-WT mice (Figure 2). However, the haplo-insufficiancy of *Brn2* (Brn2) in Braf-Pten mice significantly increased the number of tumors/mouse and the tumor growth rate (Figure 2). None of these various mouse lines generated melanomas in the absence of tamoxifen induction (except in one case of 60). None of the mice that were wild-type for Braf displayed any obvious phenotype, irrespective of the status of *Pten* or *Brn2*, including melanomagenesis and hyperpigmentation (data not shown). We verified proper defloxing of Brn2, Braf, and Pten in the melanomas (Figure S4C, D, and not shown). Thus, we compared the overall survival of human patients with a loss of one allele of PTEN who also had a loss of BRN2 (monoallelic loss) with those who had no loss of BRN2 (BRN2-normal). Patients with the loss of BRN2 showed significantly lower overall survival than BRN2-normal patients (Figure S5). In summary, our data show that Brn2 acts as a tumor suppressor *in vivo*, and its loss induces melanoma initiation and increases tumor growth rate.

**Figure 2.**
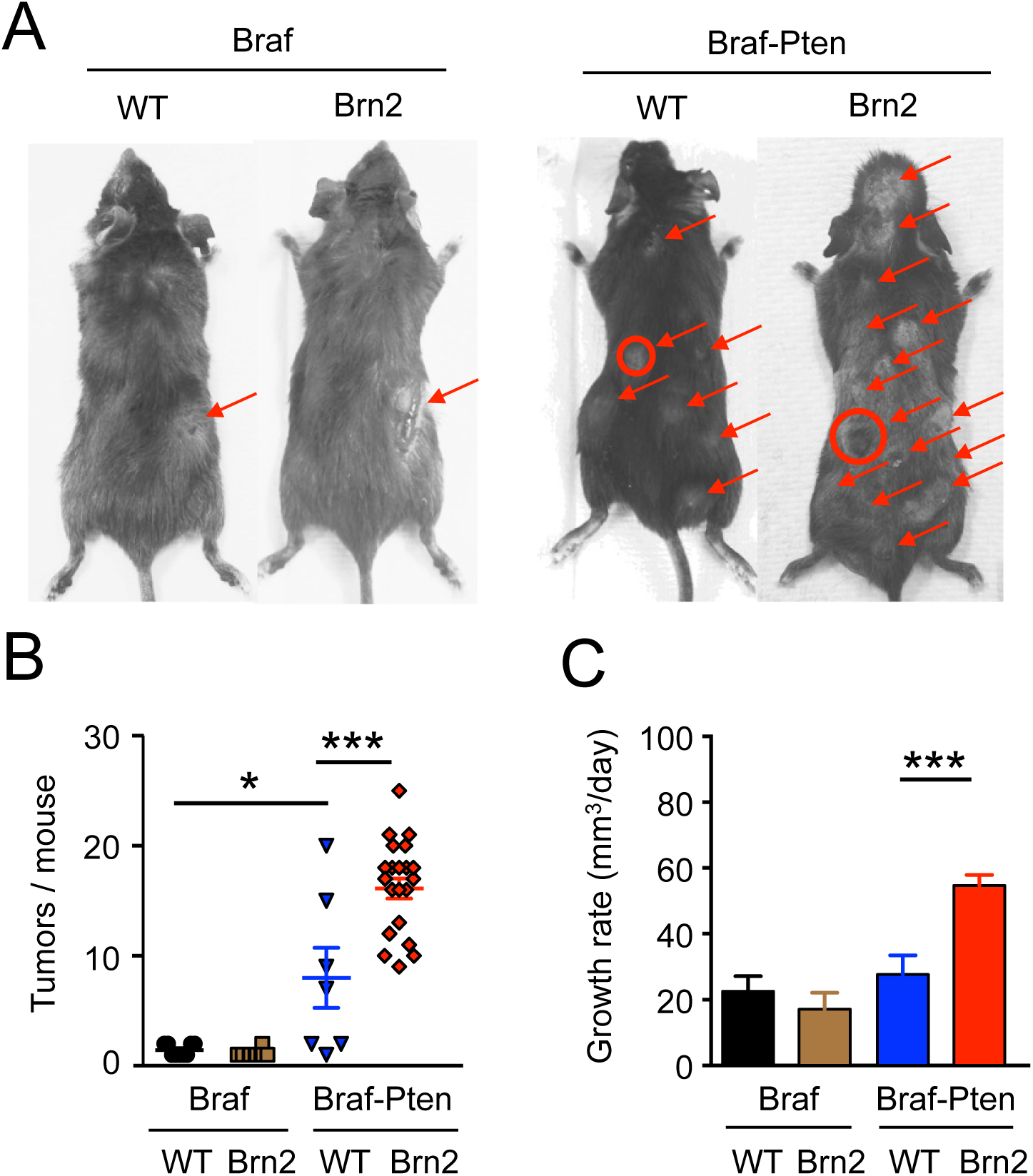
Brn2 loss potentiates melanomagenesis in Braf-Pten mice. (A) Macroscopic pictures of the dorsal view of mice with cutaneous melanomas carrying mutations in the melanocyte lineage for Braf, Pten, and Brn2 after tamoxifen induction at birth. Tyr::CreERt2; BrafV600E/+; Pten+/+ (= Braf), PtenF/+ (= Pten), Brn2+/+ (=WT), and Brn2F/+ (= Brn2). Tumors are highlighted with arrows and the sizes of the first growing tumors to appear are proportional to the diameters of the circles. (B) All Braf-WT (n = 9) and Braf-Brn2 (n = 10) mice produced cutaneous melanomas and their number was similar (1 to 2 tumors/mouse). All Braf-Pten-WT (n = 7) and Braf-Pten-Brn2 (n = 21) mice produced cutaneous melanomas. On a Braf background, in the presence of one wild-type allele of Pten (PtenF/+), the first tumors appeared after approximately 30-40 days in WT and Brn2 mice. Finally, in the absence of Pten (using PtenF/F allele; F means flox allele), the apparition of the melanoma was too rapid to observe any difference between WT and Brn2 mice (not shown). Each dot corresponds to an individual mouse. Moreover, mice were produced and not induced with tamoxifen (Braf-WT [n = 12], Braf-Brn2 [n = 25], Braf-Pten-WT [n = 7], and Braf-Pten-Brn2 [n = 13]; none of them developed melanoma after 18 months, except one Braf-Pten-Brn2 mouse that developed one melanoma after 12 months. Mice were sacrificed according to the ethical rules of the institute. (C) Growth rates of the first tumor appearing in each mouse for Braf-WT, Braf-Brn2, Braf-Pten-WT, and Braf-Pten-Brn2 mice. Statistical analysis was performed using the unpaired t-test. ns = non-significant, *p < 0.05, **p < 0.01. Error bars correspond to the s.e.m.

### Mono-allelic loss of Brn2 induces melanoma metastasis

We next evaluated the effect of Brn2 on metastasis formation *in vivo*. Since human SKCM is known to spread to proximal lymph nodes (LNs), we assessed the presence of pigmented cells in the inguinal LNs of tumor-bearing Braf-Pten mice. Specifically, we estimated the volume of the various metastasis present in LNs and the number of pigmented areas after hematoxylin & eosin (HE) staining of LN sections (Figure 3A-C). All Braf-Pten mice, irrespective of BRN2 status, showed the presence of pigmented cells in both inguinal LNs. However, the LN volume of Braf-Pten-Brn2 mice was significantly higher than that of Braf-Pten-WT (Figure 3A,B). Similarly, Braf-Pten-Brn2 LNs showed a higher number of pigmented areas per mm^2^ than Braf-Pten-WT mice (Figure 3C).

**Figure 3.**
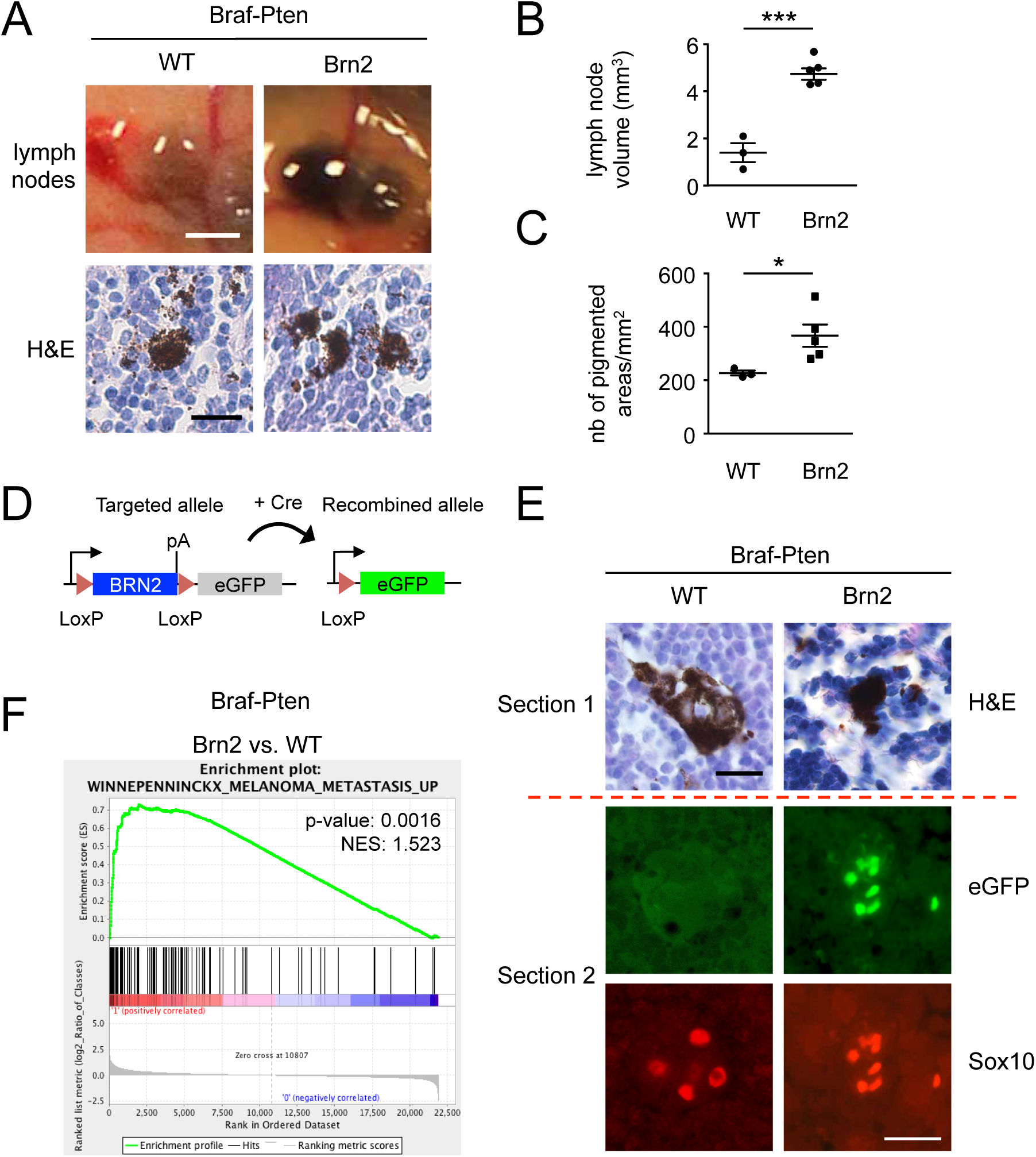
Mono-allelic loss of Brn2 induces melanoma metastasis. (A) Upper panel: Representative photomicrographs of in situ inguinal lymph nodes (LN) of Braf-Pten-WT, and Braf-PtenBrn2 mice. Scale bar = 1 mm. The pigmented volume (mm3) was estimated for each LN. Lower panel: Representative photomicrographs of hematoxylin & eosin (H&E) staining of LNs containing pigmented cells of Braf-Pten-WT, and Braf-Pten-Brn2 mice are shown. Scale bar = 20 µm. (B) Quantification of the pigmented volume of inguinal LNs in the upper panel of Figure (A). (C) Quantification of the pigmented areas per mm2 of inguinal LNs in the lower panel of Figure (A). Pigmented areas > 50 µm2 were considered. (D) Scheme showing the defloxing strategy of Brn2 in melanocytes of the primary tumor that releases eGFP expression upon the defloxing of Brn2. (E) Representative photomicrographs of serial LN sections of Braf-Pten-WT and Braf-Pten-Brn2 mice stained with H&E and the melanocyte marker Sox10. H&E staining was evaluated for one section and GFP (green channel) and Sox10 staining (red channel) evaluated for an adjacent section. Scale bar = 20 µm. (F) Gene-set enrichment analysis (GSEA) results for microarray data showing that the gene set of Braf-Pten-Brn2 (n = 5) tumors was enriched for genes involved in metastasis relative to that of Braf-Pten-WT (n = 2) tumors. Statistical analysis was performed using the unpaired t-test. Error bars correspond to the sem. ns = non-significant, *p < 0.05, ***p < 0.001, **** p < 0.0001.

To verify that the pigmented cells in the lymph nodes did not arise from cells in which the Cre recombinase had not worked efficiently, we tested whether these pigmented cells were properly defloxed for Brn2. The targeted Brn2-flox allele, used in our mouse model, has an eGFP-cassette inserted downstream of the floxed Brn2 locus (Figure 3D). Thus the production of eGFP occurs once the upstream Brn2 gene is defloxed. Consistent with correct defloxing of Brn2, pigmented areas of LNs expressed GFP in Braf-Pten-Brn2 mice, but not Braf-Pten-WT mice (Figure 3E and not shown). The pigmented cells present in Braf-Pten-WT and Braf-Pten-Brn2 LNs also expressed Sox10, a melanocytic marker (Figure 3E), that co-localized with eGFP-expression in Braf-Pten-Brn2 mice confirming the melanocytic origin of the pigmented cells observed.

To get a better understanding of the formation of metastasis in Braf-Pten-WT and Braf-Pten-Brn2, we extracted mRNA from these mouse melanomas and performed microarray-based transcriptome analysis. Gene-set enrichment analysis (GSEA) comparing Braf-Pten-Brn2 and Braf-Pten-WT melanomas revealed the former to be enriched for expression of genes shown to be upregulated in human melanoma metastases (Figure 3F).

Altogether, these data show that mono-allelic loss of Brn2 in the Braf-Pten mouse model induces LN metastasis more efficiently than Braf-Pten-WT.

### Loss of BRN2 increases cellular proliferation *in vivo* and *in vitro*

The effect of *Brn2* loss on tumor growth prompted us to investigate whether *Brn2* loss increases intra-tumoral proliferation. Staining sections for Ki-67, a marker of cycling cells, revealed that melanomas from Braf-Pten-Brn2 mice displayed a significantly higher number of Ki-67^+^ cells than Braf-Pten-WT melanomas (Figure 4A,B). To confirm this result, we injected Braf-Pten-WT and Braf-Pten-Brn2 mice with bromodeoxyuridine (BrdU) two hours prior to sacrifice, to determine whether melanoma cells were slow or fast-dividing. Braf-Pten melanomas had a significantly higher number of BrdU^+^ cells when Brn2 was lost (Figure 4C, D). These results indicated that heterozygous loss of *Pten* combined with heterozygous loss of Brn2 promotes melanoma proliferation *in vivo*. There were no detectable cells positive for cleaved caspase-3 (CCL3), an apoptosis marker, in any of these melanomas (data not shown).

**Figure 4.**
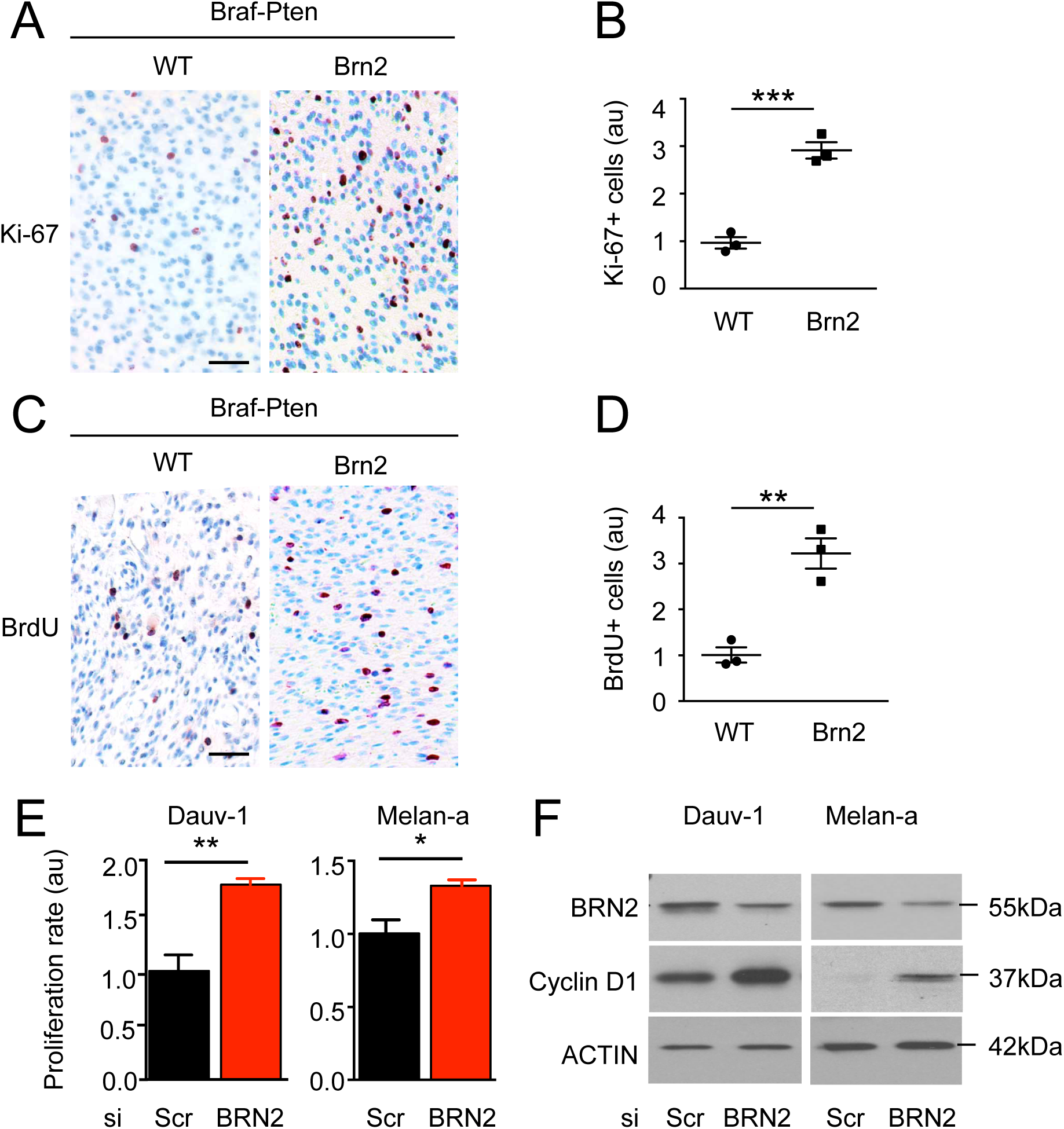
BRN2 loss induces proliferation in vitro and in vivo. (A) Representative photomicrographs of Ki-67 staining of Braf-Pten-WT and Braf-Pten-Brn2 tumors. Ki-67+ cells are stained in red. Nuclei are stained in blue. Scale bar = 40 μm. (B) Quantification of Ki-67 staining of (A). Each dot represents the result for one tumor. (C) Representative photomicrographs of BrdU staining of of Braf-Pten-WT and Braf-Pten-Brn2 tumors. BrdU+ cells are stained in red. Nuclei are stained in blue. Scale bar = 40 μm. (D) Quantification of BrdU staining of (C). (E) Growth rate of siScr and siBrn2 Dauv-1 and Melan-a cell lines (Scr = scramble). Three independent biological and technical experiments were performed for each cell line and for each condition. (F) Western blot analysis for Brn2, Cyclin D1, and actin after reduction of Brn2 in Dauv-1 and Melan-a cells. Experiments were performed independently three times. One representative blot is shown. Statistical analysis was performed using the unpaired t-test. ns = non-significant, *p < 0.05, **p < 0.01, and ***p < 0.001. Error bars correspond to the sem.

We next assessed whether Brn2 knockdown favors proliferation *in vitro* and whether this mechanism is conserved (i) between human and mouse and (ii) between “transformed” and “non-transformed” cells. We chose a human melanoma cell line, Dauv-1, and a non-transformed mouse melanocyte cell line, Melan-a. These cell lines express *Pten* and *Brn2* mRNA and protein (data not shown). Dauv-1 cells carry a BRAF^V600E/+^ mutation identical to that used in the mouse melanoma model system, and Melan-a cells are WT for Braf. siRNA-mediated knockdown of Brn2 significantly increased cell number 72 hours after transfection of both cell lines (Figure 4E, Figure S6E). Brn2 knockdown, assessed by western blot, led to an increase in cyclin D1 protein levels in both cell lines, but did not alter cyclin D1 mRNA levels, suggesting regulation of cyclin D1 at the protein level (Figure 4F, Figure S6F, G). Overall, Brn2 loss induces the proliferation of cells of the melanocytic lineage *in vivo* and *in vitro*.

### Brn2 loss in Braf-Pten melanoma drives cellular proliferation through the PI3K-pathway

The results presented so far suggest that monoallelic loss of *Brn2* promotes melanoma formation and proliferation. To achieve this biological effect, presumably Brn2 must affect pro-proliferative signals and downstream transcriptional programs. To investigate the mechanism underlying the effect of Brn2-loss we extracted mRNA from Braf-Pten-WT and Braf-Pten-Brn2 mouse melanomas and performed microarray-based transcriptome analysis. We detected 290 differentially expressed genes between Braf-Pten-Brn2 and Braf-Pten-WT tumors, of which 141 were downregulated and 149 upregulated (log2 (fold change) > +1 and log2 (fold change) < −1, with a significance threshold: p < 0.05, Table S3). Gene ontology analysis for cellular processes (GO Cellular process 2018) showed pathways associated with cellular proliferation (including cell-cycle transition, mitotic spindle formation, and chromosome segregation) to be upregulated by Brn2 heterozygous loss in Braf-Pten melanomas (Figure 5A). In terms of molecular function (GO Molecular function 2018), several microtubule and kinase-associated processes were up-regulated, consistent with the induction of cellular proliferation. These data were supported by GSEA, showing that gene signatures related to proliferation and cell-cycle scores were enriched in Braf-Pten-Brn2 melanomas in comparison to Braf-Pten-WT melanomas (Figure 5B). We next assessed the signaling pathways upregulated in Brn2-deficient melanomas. Enriched pathway analysis (WikiPathways 2016) revealed the upregulation of pathways that affect the cell cycle, DNA replication, the G1/S transition, DNA damage response and retinoblastoma (RB)-signaling (Figure 5C). KEGG pathway analysis (KEGG 2016) also showed the upregulation of cell-cycle and DNA-replication signaling, as well as the activation of PI3K-Akt signaling (Figure 5D). Significantly, activation of PI3K signaling was verified by performing western blots of Braf-Pten tumor samples for members of the PI3K-AKT pathway, with Braf-Pten-Brn2 melanomas showing higher levels of pAkt and pS6 than Braf-Pten-WT tumors (Figure 5E). Overall, these results show that Brn2 loss in Braf-Pten melanomas can drive cellular proliferation and co-occurs with activation of the PI3K-AKT pathway.

**Figure 5.**
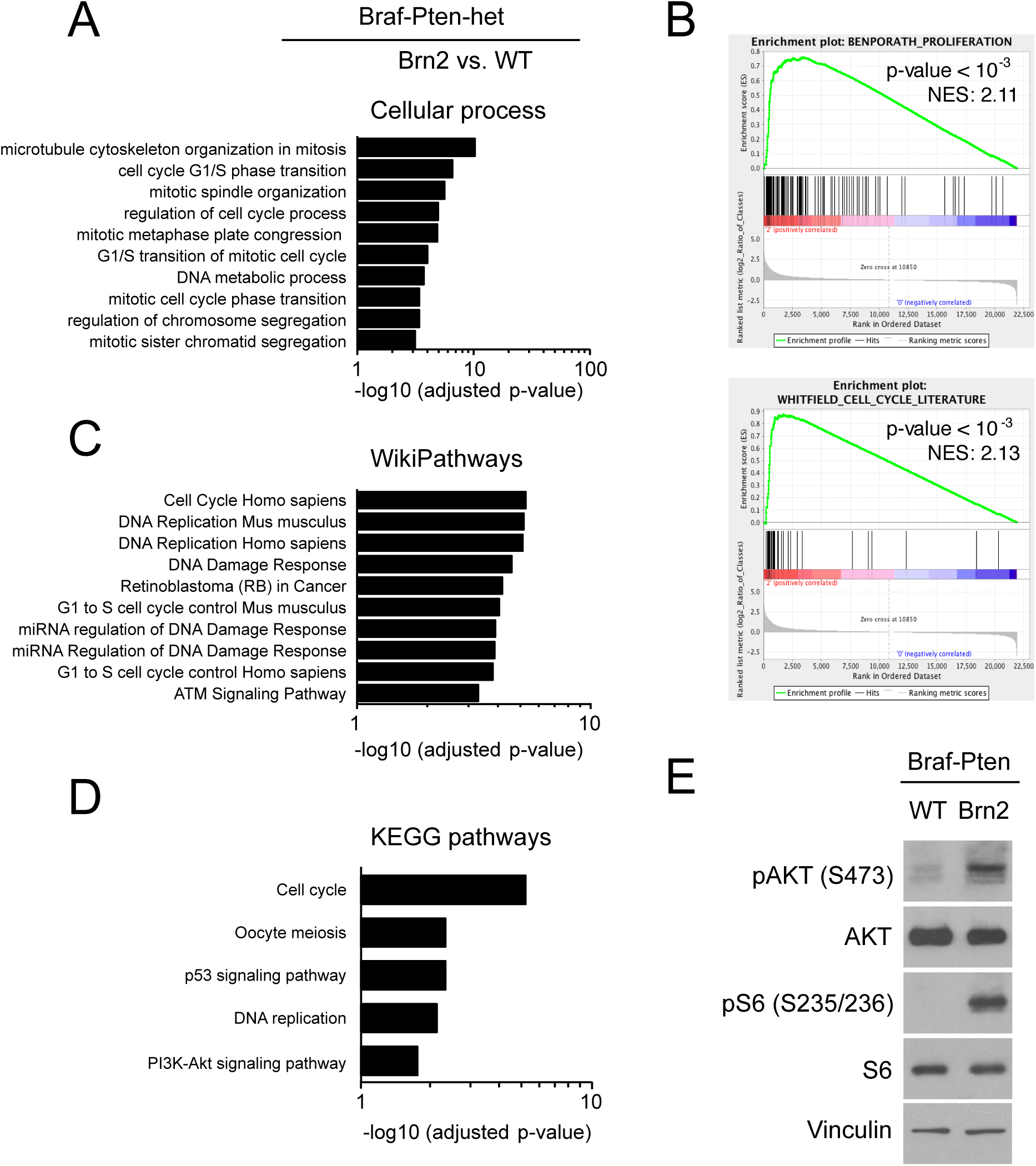
Brn2 loss in Braf-Pten melanoma activates cellular proliferation through the PI3K-Akt pathway. (A) Gene ontology analysis of cellular processes (GO Cellular Process 2018) for microarray data of total RNA extracted from Braf-Pten-Brn2 (n = 6) vs. that from Braf-Pten-WT (n = 2) mouse melanoma. (B) Gene-set enrichment analysis (GSEA) of Braf-Pten-Brn2 and Braf-Pten-WT. Enriched pathway analysis (C) WikiPathways 2016, and (D) KEGG 2016 for microarray data of total RNA extracted from Braf-Pten-Brn2 and Braf-Pten-WT melanoma. (E) Western blot results for phospho-AKT, Akt, phospho-S6, S6 and vinculin in Braf-Pten-WT and Braf-Pten-Brn2 melanoma. For each graph, all hits were ranked according to the adjusted p-value from low (top) to high (bottom). An adjusted p-value of 0.05 was used as a threshold for all gene-ontology and enriched pathway analysis.

### BRN2 binds to the *PTEN* promoter and BRN2 loss leads to the reduction of *PTEN* transcription

The PI3K-AKT pathway is induced in melanoma and its induction abrogates BRAF^V600E^-induced senescence ^32,33^. The loss of *Pten*, a suppressor of the PI3K pathway, induces melanoma initiation and proliferation *in vivo* ^3,30^. Since our Braf-Pten mouse melanoma model retained one functional allele of *Pten*, we hypothesized that *Brn2* loss would induce a reduction in the level of expression from the WT *Pten* allele, leading to increased PI3K-AKT signaling and consequent melanoma initiation and proliferation. We therefore evaluated Pten protein levels in Braf-Pten tumors by immunohistochemistry and found that Braf-Pten-Brn2 tumors showed fewer Pten^+^ cells than Braf-Pten-WT tumors (Figure 6A). This result was verified by western blotting of Braf-Pten tumor samples. The reduction of Brn2 correlates with the reduction of Pten in Braf-Pten-Brn2 tumors compared to the Braf-Pten-WT tumors (Figure 6B). In accordance with the protein levels, the mRNA levels of *Brn2* and *Pten* were significantly lower in Braf-Pten-Brn2 tumors than in Braf-Pten-WT tumors, suggesting regulation of *Pten* at the transcriptional level (Figure 6C).

**Figure 6.**
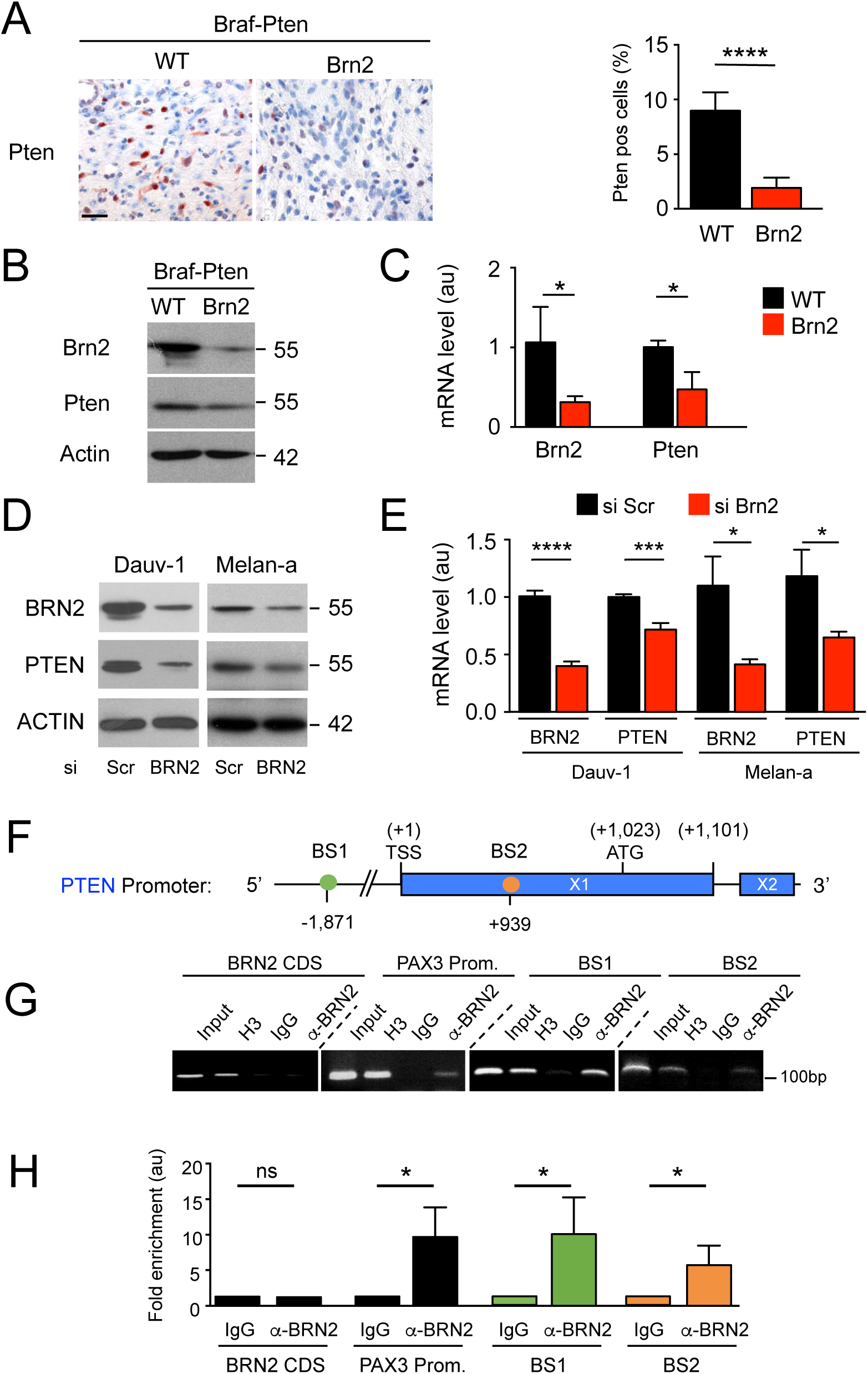
Brn2 binds to the Pten promoter and Brn2 loss leads to Pten transcription reduction. **(A)** Representative photomicrographs of immunohistochemistry staining of Pten (red) in Braf-Pten-WT and Braf-Pten-Brn2 mouse melanomas are shown. Scale bar = 40 µm. The percentage of Pten^+^ cells in WT and mutant tumors is shown. **(B)** Western blot analysis of Brn2, Pten and actin for Braf-Pten-WT and Braf-Pten-Brn2. One representative example is presented. **(C)** RT-qPCR of Brn2 and Pten from Braf-Pten-WT and Braf-Pten-Brn2 melanomas. Three independent mouse melanomas per genotype were analyzed. Data were normalized against the values of Gapdh. au = arbitrary unit. **(D)** Western blot analysis of BRN2, PTEN and ACTIN from Dauv-1 human melanoma cells and Melan-a mouse melanocytes after siRNA mediated knockdown. A representative western blot is shown. Scr = Scramble. **(E)** RT-qPCR of *BRN2* and *PTEN* from human melanoma cells (Dauv-1) and mouse melanocytes (Melan-a) after siRNA-mediated knockdown. Specific primers were used for human and mouse samples. Dauv-1 (n = 6), Melan-a (n = 4). Data were normalized against the values for *TBP* (Dauv-1) or *Gapdh* (Melan-a). **(F)** Scheme of the human PTEN promoter containing two BRN2 binding sites (BS) represented as colored circles. Note that BS are conserved between humans and mice. TSS = transcription start site. Exons (X) 1 and 2 are shown as horizontal rectangles. The translation start site (ATG) and the end of exon 1 are indicated. All numbering is relative to the TSS (+1). **(G)** ChIP assays of BRN2 binding to the PTEN promoter in Dauv-1 melanoma cells. All data shown are representative of at least three independent assays. **(H)** Quantification of the ChIP-qPCR, plotted and normalized against IgG as the reference. au = arbitrary unit. Statistical analysis was performed using the unpaired t-test. Error bars correspond to the sem. ns = non-significant, *p < 0.05, ***p < 0.001, and ****p < 0.0001.

Next, we verified the mechanism of Pten repression mediated by reduced levels of Brn2 in the cell lines Dauv-1 and Melan-a. siRNA-mediated BRN2 knockdown led to significantly reduced PTEN protein and mRNA levels in these cell lines (Figure 6D,E and Figure S7A). We examined the *PTEN* promoter for BRN2 binding sites conserved between humans and mice to determine whether BRN2 acts directly on *PTEN* and detected two Brn2 binding sites (BS1 and BS2) at positions − 1,871 and +939 (numbering relative to ATG) (Figure 6F). Chromatin-immunoprecipitation (ChIP) for BRN2, performed on Dauv-1 melanoma cells extracts, followed by qPCR, revealed quantitative BRN2 binding to both binding sites, comparable to BRN2 binding on the *PAX3* promoter (Figure 6G,H).

The *MITF* gene encodes a key transcription factor that plays a major role in melanocyte and melanoma biology (Goding and Arnheiter, submitted review). Several studies have reported that *MITF* transcription is directly repressed by BRN2 whereas a reduction in BRN2 levels leads to increased MITF levels ^12,19,34^ and a study showed that BRN2 induces MITF level ^35^. We therefore examined whether BRN2 loss can also repress PTEN indirectly via the induction of MITF. siRNA-mediated *MITF* knockdown in three human melanoma cell lines (501mel, SK28, HBL) led to a significant increase of PTEN protein and mRNA levels (Figure S8A, B). Whole genome ChIP-sequencing revealed that MITF binds to an E/M-box 3’ of the human *PTEN* gene (Figure S8C, D). Quantitative ChIP experiments confirmed binding of MITF to this specific site that was greater than at the TYR promoter, a well-characterized MITF target (Figure S8E, F). Overall, the reduction of MITF levels induces an increase in the level of PTEN mRNA. As such, it is plausible that Brn2 induces *Pten* transcription through two different mechanisms, but concurrent mechanisms: (i) directly through BRN2 binding to the *PTEN* promoter to induce its transcription and (ii) indirectly via BRN2 repressing MITF expression, with MITF binding to the 3’ end of PTEN to inhibit its transcription. In the absence of BRN2, these mechanisms are disrupted and PTEN transcription is downregulated.

Overall, BRN2 loss reduces PTEN transcription *in vitro* and *in vivo*, thus ramping up PI3K signaling and inducing both the initiation of melanoma and the formation of metastases.

## DISCUSSION

A well-established principle of cancer biology is that tumors are initiated by a combination of oncogene activation together with loss of tumor suppressor expression or activity. In melanoma key oncogenic drivers, such as BRAF and NRAS, have been well defined. Loss of P16 or PTEN tumor suppressor activity is required to bypass oncogene-induced senescence and permit melanoma initiation. However, while inactivation of tumor suppressors by mutation has been extensively studied, less well understood is how their activity may be modulated by changes in their mRNA expression mediated by key melanoma-associated transcription factors. Here, we identify BRN2, a key transcription factor lying downstream of three melanoma-associated signaling pathways (WNT/β-catenin, MAPK, and PI3K), as a novel tumor suppressor that functions to regulate PTEN expression. Thus monoallelic loss of BRN2 promotes melanoma initiation in a Braf^V600E^/Pten^+/-^ background where mono-allelic loss of *Pten* sensitizes cells to loss of *Brn2*.

Previous work has primarily linked BRN2 to melanoma invasion *in vitro* and in *in vivo* xenograft experiments ^12,36^, but its role during melanoma initiation and proliferation *in vivo* and in normal melanocytes had not been determined. We report that, consistent with *BRN2* playing a key role as a tumor suppressor in melanomagenesis, its locus is frequently lost in human skin cutaneous melanoma (SKCM) metastases, independently of their NRAS or BRAF status, and that overall patient survival is dependent on their BRN2 status. Significantly, the overall survival of patients with a mono-allelic loss of PTEN is higher when the *BRN2* locus is intact.

Although these observations are consistent with BRN2 affecting human melanoma initiation and progression, loss of the BRN2 locus is frequently associated with large segmental deletions that affect the long arm of chromosome 6 (6q). Moreover, *in vitro* studies have shown that several genes are linked to melanomagenesis in the co-deleted region, including *ARID1B, MCHR2, CCNC, CDK19, DLL1, ROS1*, and *CRYBG1/AIM1*, although none have yet been shown to be functionally important in melanoma *in vivo* ^34,37-43^.

Thus it might be argued that any of the genes located in this frequently deleted region may be acting to modulate melanoma initiation or progression. However, our functional mouse molecular genetics models show conclusively that the heterozygous loss of Brn2 promotes melanoma initiation and initial growth. Similarly, our transcriptomic analysis showed that the mRNA levels of the seven genes in the region co-deleted in humans (*Arid1b, Mchr2, Ccnc, Cdk19, Dll1, Ros1*, and *Crybg1/Aim1*) were not affected in the mouse tumors (Braf-Pten-WT and Braf-Pten-Brn2). Collectively, these results strongly suggest that the reduction of Brn2 levels is the critical event that cooperates with heterozygous Pten to promote the initiation and growth of melanoma, independently of the co-deleted genes in melanoma. Our observations are therefore consistent with BRN2 acting as a novel tumor suppressor in melanoma, and are in full agreement with the predominantly mutually exclusive pattern of BRN2 and Ki-67 *in situ* staining of invasive melanoma ^20^.

The presence of Braf^V600E^ promotes proliferation prior to inducing senescence and the loss of Pten results in senescence bypass ^3,27,30^. As such, we believe that the increased proliferation and tumor-initiation frequency observed in our Braf^V600E^/Pten^+/-^ model arising as a consequence of the reduction of Brn2, is likely to occur as a consequence of the ability of Brn2 to activate Pten expression and suppress PI3K signaling either directly or indirectly via Mitf repression. In other words, inactivation of Brn2 or a reduction in its expression would lead to low expression of the remaining Pten allele and as a consequence increase the probability of senescence bypass. Our *in vivo* data therefore reveal that melanocyte-specific Brn2 deletion in Braf-Pten mice promotes the initiation of melanoma and establish Brn2 as a central tumor suppressor, acting at different steps of melanomagenesis, and complement numerous other studies showing the effect of Brn2 on invasion ^12-14,20,21,36^.

Brn2 heterozygous mice were more prone to form LN metastases efficiently than mice that were Brn2 WT when Pten was already heterozygous. The efficiency of the metastasis process depends on the status of Brn2 as the number and size of each micro-metastasis was greater in Brn2 heterozygous than Brn2 wild type melanoma. This is important as mono-allelic loss of *BRN2* in human, corresponding to *BRN2* heterozygous melanoma, occurs in 53% of human melanoma. In Brn2 wild-type melanoma, the change of the level/activity of Brn2 is possible but the amount of protein found in Brn2 wild-type melanoma absorbs better transient Brn2 depletion than in Brn2 heterozygous melanoma.

Some residual BRN2 activity might be required for efficient melanoma progression. The formation of metastases is a multistep process in which cells proliferating in the primary tumor, and surviving the metastatic process, must undergo a switch to an invasive phenotype prior to a switch to a proliferative phenotype on site. Since the switch from proliferation to invasion, and back, has been associated with the activity of MITF, it is possible that for efficient metastatic colonization cells must be able to modulate MITF expression via BRN2. In this respect, it is especially important to note that BRN2 has been identified as a key regulator of MITF ^12,35^. Consistent with the observation that MITF and BRN2 are frequently observed in mutually exclusive populations in melanoma, and that BRN2 may act *in vivo* as an MITF repressor ^12^, we have observed that in a non-tumoral context, the specific knock-out of Brn2 *in vivo* in melanocytes increases the level of Mitf (publication in preparation). Importantly, during progression of mouse Braf-Pten melanoma Mitf levels are modulated. During the initial phase of growth, melanoma cells are pigmented, indicative of Mitf activity, but later *in situ* they lose pigmentation and ability to produce Mitf (Laurette *et al*. submitted, and not shown). Although the reduction of Brn2 in Braf-Pten-Brn2 primary melanoma is not sufficient to re-induce Mitf mRNA and pigmentation in all cells of these primary melanomas, the Braf-Pten melanoma cells that formed LN metastases were pigmented and re-expressed Sox10, a key transcription activator of Mitf. Thus, during progression of Braf-Pten melanomas Mitf is produced during initial growth and subsequently repressed during the second phase, before being re-expressed in LN metastases. It seems therefore likely that one role of the residual BRN2 in the heterozygotes may be to facilitate the modulation of MITF expression during metastatic spread.

Although here we have focused on the role of BRN2 in melanoma, BRN2 is also expressed in a number of other cancer types including small cell lung cancer, neuroblastoma, glioblastoma and neuroendocrine prostate cancer ^8-10^. While BRN2-mediated regulation of MITF is not likely to be important for non-melanoma cancers that do not express MITF, the ability of BRN2 to modulate PTEN expression uncovered here may play an equally important role in promoting the initiation and progression of these cancer types. In this respect the inducible knockout mice described here may represent an important tool to examine the role of BRN2 in non-melanoma cancers.

In conclusion, our results identify Brn2 as a key tumor suppressor through its ability to modulate Pten expression that, given the high prevalence of monoallelic mutations, is likely to play a key role in initiation of human melanoma and likely other BRN2-expressing cancer types. Since BRN2 expression is activated by PI3K signaling via PAX3 ^15^, its ability to suppress PI3K signaling by increasing PTEN expression may also provide cells with a negative feedback loop to control the PI3K pathway. Moreover activation of BRN2 by MAPK signaling downstream BRAF ^14^ as well as WNT/β-catenin signaling ^13^, may also permit coordination between these pathways and the PTEN/PI3K axis. Finally, given the importance of BRN2 in melanomagenesis identified here as well as its frequent heterozygosity, it may be important to determine whether tumors with low BRN2 expression may be more susceptible to PI3K pathway inhibition. In this respect our newly developed mouse model of BRN2-deficient melanoma could be useful for the pre-clinical testing of inhibitors for clinical development especially since it has been shown that BRN2 is involved in DNA repair ^44^.

## ONLINE METHODS

Detailed materials and methods are given in the supplemental information.

### TCGA data mining

All TCGA data sets for somatic mutations, copy number alterations (CNAs), RNA levels, and clinical data for skin cutaneous melanoma and other cancers were retrieved from http://www.cbioportal.org on May 28, 2017.

### Copy number analysis

Copy number data was obtained from a previous study, prior to be evaluated, calibrated and segmented according to ^46-48^. A cut-off of −0.3 was set to determine mono-allelic loss of BRN2. Samples with GISTIC copy-number values of “-1” were considered as mono-allelic loss (red) and samples with GISTIC copy-number values of “-2” as bi-allelic loss. TCGA dataset CNA-data set (n = 367). BRN2 CNA gain/amplification (GISTIC > “+ 1” or “+2”) not shown (n = 29, 7.9%). Known genes linked to melanomagenesis are indicated at their respective location. Vertical lines indicate the extent of genetic deletion associated with *BRN2* loss in individual patients. Nine patients with bi-allelic loss (orange) and ten randomly chosen patients with mono-allelic loss (red) are shown. No data for the deleted region were available for one patient with bi-allelic loss. Deleted genomic regions of each patient were based on the TCGA CNA-data set (n = 367). Patient IDs with the exact extent of genetic deletion can be found in Table S2.

### Level of expression of BRN2 mRNA in melanoma patients

The correlation between BRN2 mRNA levels and Clark levels of SKCM samples was evaluated from a total of 316 independent melanoma including Level II, (n = 18), Level III (n = 77), Level IV (n = 168), Level V (n = 53). Data for Level I (n = 6) are not shown. BRN2 mRNA levels were calculated from RNA sequencing read counts using RNA Seq V2 RSEM and normalized to transcripts per million reads (TPM). The means are shown.

The correlation between BRN2 mRNA levels and Breslow index in SKCM samples was evaluated from a total of 210 independent melanoma including Stage 1 (< 0.75 mm, n = 26), Stage 2 (0.75-1.50 mm, n =53), Stage 3 (1.51-2.25 mm, n = 28), Stage 4 (2.26-3.00 mm, n = 26), and Stage 5 (> 3mm, n = 77). BRN2 mRNA levels were calculated from RNA sequencing read counts using RNA-Seq V2 RSEM and normalized to transcripts per million reads (TPM). The means are shown.

Kaplan-Meier curves comparing 30-year overall survival of SKCM patients to BRN2 mRNA levels were established as followed: BRN2 mRNA levels were calculated from RNA Sequencing read counts using RNA-Seq V2 RSEM and normalized to transcripts per million reads (TPM) based on the TCGA melanoma metastasis data set (total n = 359): BRN2 High (> 1 TPM, black, n = 133) and BRN2 low (≤ 1 TPM, red, n = 226).

### Microarray analysis

11μg cRNA were hybridized to mouse MOE430 gene expression Affymetrix microarrays (Affymetrix, #900443), washed and stained (Affymetrix fluidics station 450), and scanned (Affymetrix GeneChip Scanner 3000). Normalization and evaluation of the expression were performed using the PLIER and edgeR packages. Gene-ontology and enriched pathway analysis was performed using Enrichr ^45^. Gene-set enrichment analysis (GSEA) was performed using gene sets from the Molecular Signatures Database (MSigDB v6.2). Microarray-based transcriptomic analyses of total RNA (RIN > 7) extracted from tumors (n = 13) were subjected to several normalization steps. The threshold of differentially expressed genes from Braf-Pten-Brn2 *vs*. Braf-Pten-WT tumors was defined as logFC > +1 and > −1 with p > 0.05. Gene-ontology and enriched pathway analysis of 290 differentially expressed genes (141 down- and 149 upregulated genes) was performed using Enrichr ^45^.

### Mouse Models

Mice were bred and maintained in the specific pathogen-free mouse colony of the Institut Curie, in accordance with the institute’s regulations and French and European Union laws. The transgenic *Tyr::Cre*^*ERT2*^, *Braf*^*V600E/+*^, *Pten*, and *Brn2* mice have been described, characterized, and backcrossed onto a C57BL/6 background for more than ten generations ^26-28,31^. Genotyping were performed accordingly (Tables S3 and S4)

### In vivo gene activation/deletion and melanoma monitoring

Newborn mice were treated dorsally with 0.4μg/day/mouse tamoxifen from P1 to P3.. Non-tamoxifen-induced mice of the same genotype were used as controls. Developing skin excrescences > 3 mm diameter were considered to be melanomas, and validated after growth. Mice were sacrificed and autopsied four weeks after tumor appearance or once the tumors reached 2 cm^3^. Mouse melanomas were excised, rinsed in cold PBS snap-frozen in liquid nitrogen for subsequent transcriptomic and western blot analysis and fixed in 4% PFA and embedded in paraffin or OCT for histological analysis and immunostaining.

### Immunohistochemistry of mouse melanoma and inguinal lymph nodes

Paraffin-embedded mouse melanomas were sectioned into 7-μm-thick transverse sections and stained with hematoxylin/eosin (H&E), as previously described ^49^. Antibodies are from Nova-Costra for Ki-67, BD Biosciences for BrdU, from Abcam for SOX10 and Cell signaling for PTEN. Images were captured using a ZEISS Axio Imager 2 with Axiocam 506 color cameras. Image analysis was performed using ZEISS ZEN, Adobe Photoshop, and *ImageJ* software. Quantifications of Ki-67 and BrdU stainings were determined as a percentage. The percentage of Ki-67^+^ and BrdU^+^ cells from three fields (1,000-2,000 cells/field) from three independent tumors per genotype was determined and normalized.

### Protein extraction from tumors

All steps were performed at 4°C. Tissues were transferred to a tube containing 2.8-mm stainless steel beads and 1mL RIPA buffer supplemented with sodium orthovanadate prior adding complete protease inhibitor and Phostop (Roche). Tissues were homogenized followed by centrifugation at 13,000 rpm for 1 min. Supernatants were transferred and centrifuged for 20 min at 12,000 rpm. Supernatants were collected and incubated with 200µL previously PBS-washed G-Sepharose beads for 2h. Samples were centrifuged at 12,000 rpm for 5 min and quantified using the Bradford assay.

### Cell lines

Melan-a and C57BL/6 9v cells were grown as previously described ^4,50^. 501mel, 501mel-MITF-HA, HBL, SK-Mel-28, and Dauv-1 cells were grown in RPMI 1640 media supplemented with 10% FCS and 1% Penicillin-Streptomycin ^51-53^. Cells are routinely tested for the absence of mycoplasmas using MycoAlert (Lonza). The origin of the cell lines is given in the supplemental information.

### Genomic DNA extraction from cell culture

Genomic DNA was extracted using the AllPrep DNAMini Kit (Qiagen) according to manufacturer’s instructions (Tables S4 and S5).

### siRNA-mediated knock-down

siRNA targeting human BRN2 and MITF was purchased from Dharmacon as a SMART-pool mix of four sequences. siRNA targeting PTEN from Santa Cruz Biotech. Si Scramble (siSCR), with no known human or mouse targets, was purchased from Eurofins Genomics (Table S6). Briefly, cells were transfected with Lipofectamin2000 with 200 pmol siRNA or siSCR and assayed for mRNA expression or protein content 48 or 72 h post-transfection.

### Western blotting and detection

Whole-cell lysate was prepared from cell lines using RIPA buffer supplemented with sodium orthovanadate, complete inhibitor, and Phostop. SDS-PAGE was carried out on 10% polyacrylamide protein gels. Following the transfer of the proteins, the nitrocellulose membranes were blocked in TBST with 5% non-fat dry milk for 1.5h at RT and then probed with various antibodies. BRN2, Cyclin D1, PTEN, phospho-S6, S6, phospho-AKT, and AKT antibodies were from Cell signaling, MITF from Abcam, β-actin and inculin from Sigma. Secondary antibodies were either HRP-conjugated goat anti-rabbit or anti-mouse IgG. Blots were incubated in ECL (Pierce) and revealed using ECL hyperfilm (GE Healthcare). All primary antibodies were used at a dilution of 1/1,000, except β-actin and vinculin (1/5,000). All secondary antibodies were used at a dilution of 1/20,000. Quantification of the western blots was performed using *ImageJ* software.

### Chromatin immunoprecipitation

ChIP experiments were performed as previously described ^4^. ChIP assays of BRN2 binding to the PTEN promoter. ChIP assays were performed using an antibody against BRN2 and analyzed after 30-cycle PCR in exponentially growing phase of Dauv-1 melanoma cells. PAX3 promoter (prom.) and Brn2 coding sequences (CDS) were used as positive and negative controls, respectively. Input represents approximately 0.4% of the input used for the ChIP assay. H3 (histone H3) and IgG (Immunoglobulin G) were used as positive and negative controls for each region of interest, respectively. The oligonucleotides, their position on the genome, and the sizes of the amplified fragments are given in Tables S4 and S5.

### RNA extraction and (ChIP) RT-qPCR

Tissues were crushed with a mortar and pistol, and stainless steel beads. Qiazol was used to homogenize the samples prior extracting RNA using the miRNeasy Kit. Purified RNA was reversed transcribed using M-MLV Reverse Transcriptase. Real-time quantitative PCR (qPCR) was performed using iTaq™ Universal SYBR Green Supermix. Each sample was run in technical triplicates and the quantified RNA normalized against TBP (human) or Gapdh (mouse) as housekeeping transcripts (Tables S4, S5, and S7)

### CHIP-SEQ

Chip-seq analysis was performed as mentioned in the study by the Goding lab (submitted for publication – see supplemental information)

### Softwares

GraphPad PRISM, Adobe Illustrator, Adobe Photoshop, and Microsoft Power Point software were used to generate all graphs and figures.

### Quantification and statistical analysis

Cell culture-based experiments were performed in at least biological triplicate and validated three times as technical triplicates. P-values for the comparison of two groups were calculated using the unpaired Student t-test or Mann-Whitney test. P-values for the comparison of multiple groups were calculated using the analysis of variance (ANOVA) and Fisher’s least significant difference tests. P-values for categorical data were calculated using the Chi-square test. P-values for the comparison of Kaplan–Meier curves were calculated using the log-rank (Mantel–Cox) or Gehan-Breslow-Wilcoxon test giving weight to the early events. Exact p-values were calculated as reported by Prism 6.

### Data and software availability

Data Resources. The accession number for the sequencing data reported in this paper is dbGaP: phs001550.v1.p1.

Mouse melanoma tumors microarrays has been deposited on Gene Expression Omnibus (GEO) website under accession number GSE126524 https://www.ncbi.nlm.nih.gov/geo/query/acc.cgi?acc=GSE126524

aCGH information of the human melanoma cell lines are presented as an excel file (Table S1 aCGH human melanoma cell lines.xlsx).

## Acknowledgements

We are grateful to Dorothy C Bennett, Ghanem Ghanem, Florence Faure, and Meenhard Herlyn for providing cell lines. Hong Wu for providing Pten flox mice. We thank the Institut Curie staff responsible for the animal colony (especially P. Dubreuil), and the histology (S. Leboucher), FACS (C. Lasgi), and PICT-IBiSA imaging (C. Lovo) facilities. This work was supported by the Ligue nationale contre le cancer, INCa, ITMO Cancer, Fondation ARC (PGA), and is under the program «Investissements d’Avenir» launched by the French Government and implemented by ANR Labex CelTisPhyBio (ANR-11-LBX-0038 and ANR-10-IDEX-0001-02 PSL). MH had a fellowship from PSL and FRM, PS had a fellowship from INSERM, and MLC had a fellowship from FRM. CRG was supported by the Ludwig Institute for Cancer Research. RC was supported in part by a grant from the National Institutes of Health (AR062547, RAC) and a postdoctoral fellowship from the American Association for Anatomy (CK). ES was supported from the Research Fund of Iceland (184861-053). LL and ES are supported by a Jules Verne grant.

---

**Figure S1.**
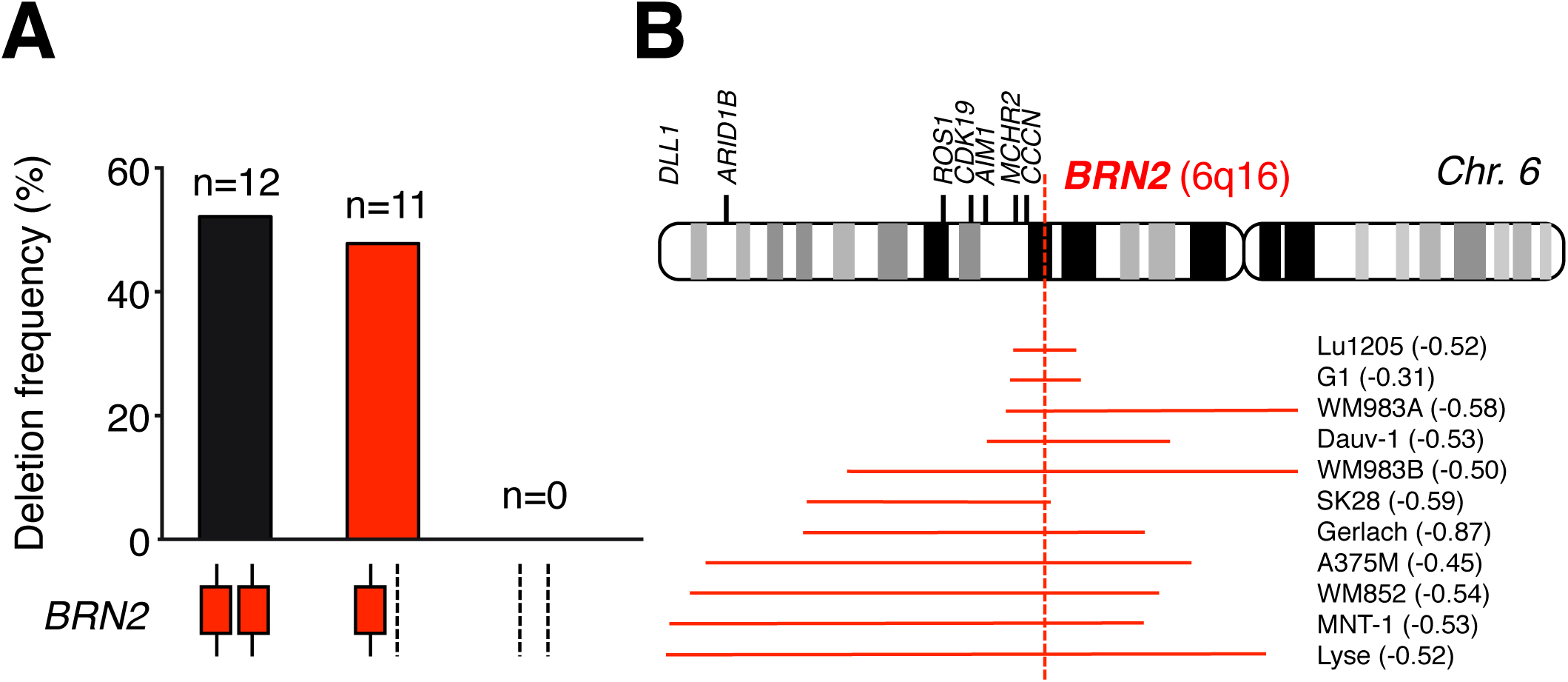
BRN2 is frequently affected by large segmental deletions in human melanoma cell lines. (A) Bar graphs showing the frequency of copy-number alterations (CNAs) of BRN2 in human melanoma cell lines (n = 23). CNAs were inferred from the log2 ratio of comparative genomic hybridization for each cell line: log2 = 0 was considered as diploid (black), −0.3 > log2 > −1 as mono-allelic loss (red), and log2 ≤ −1 as bi-allelic loss. CNAs of BRN2 are indicated in the pictograms (red rectangles and dashed lines) under the graphs. (B) Pictogram showing the extent of segmental deletion affecting the BRN2 locus on Chr.6q16 (dashed red line) in human melanoma cell lines. Known genes linked to melanomagenesis are indicated at their respective location. Horizontal lines indicate the extent of genetic deletion associated with BRN2 loss for each cell line (red). The name of the respective cell line and the log2 ratio from the comparative genomic hybridization is indicated.

**Figure S2.**
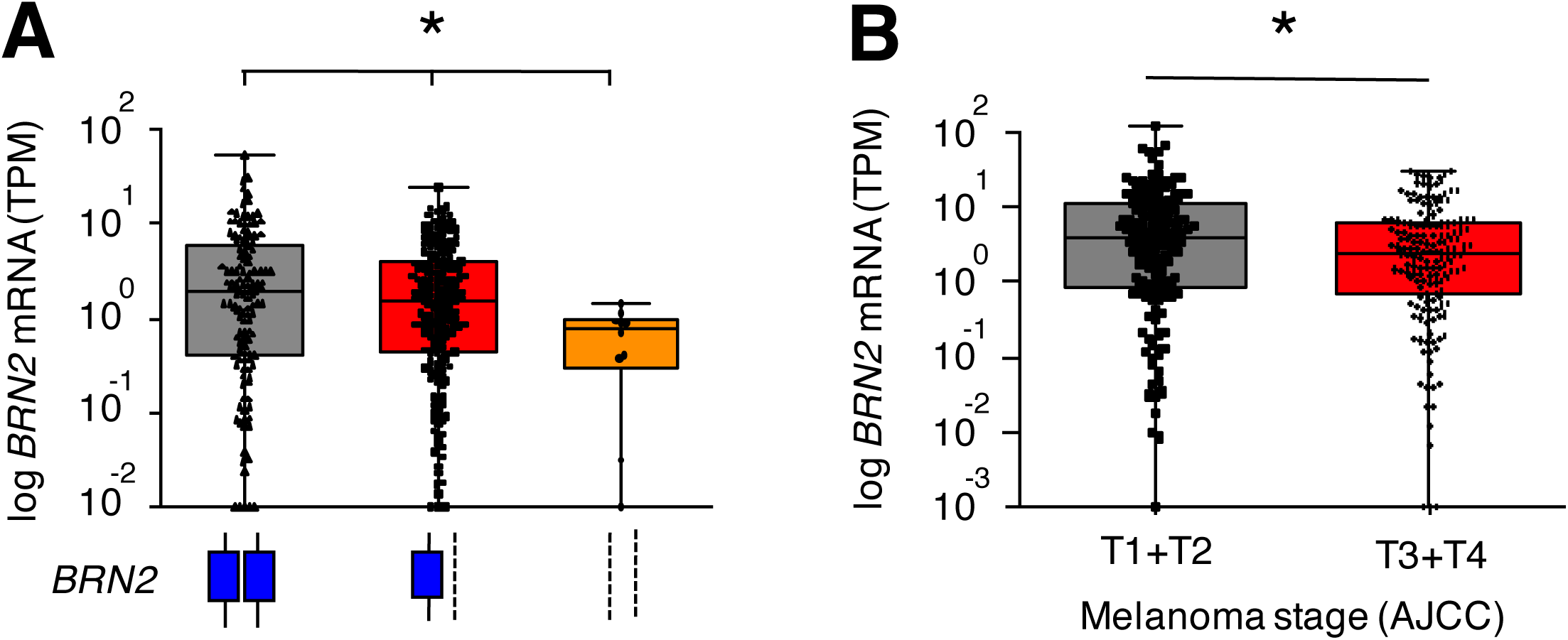
BRN2 loss is associated with lower BRN2 mRNA levels and higher AJCC Stage. (A) Box and whisker plot of the correlation between BRN2 mRNA levels and BRN2 CNAs in human SKCM metastases. BRN2 mRNA was normalized to transcripts per million (TPM). BRN2 CNAs were estimated from GISTIC values. Diploid = “0”, monoallelic loss = “-1”, bi-allelic loss = “-2”, gain = “+1”, and high-level amplification = “+2”. BRN2 bi-allelic loss (n = 10), BRN2 monoallelic loss (n = 195), and BRN2 diploid (n = 133). BRN2 Gains and high-level amplification (n = 27 + 2) are not shown. TCGA-CNA data set (n = 367). Data are represented as the median ± interquartile range (box) and ± 100% range (whiskers). (B) Box and whisker plot of the correlation between BRN2 mRNA level and melanoma stage (AJCC, T1-T4). Stages T1 and T2 were combined into one group and T3 and T4 into another. Error bars correspond to the sem. Statistical analysis were performed using ANOVA (A) or Mann-Whitney (B) tests, *p < 0.05.

**Figure S3.**
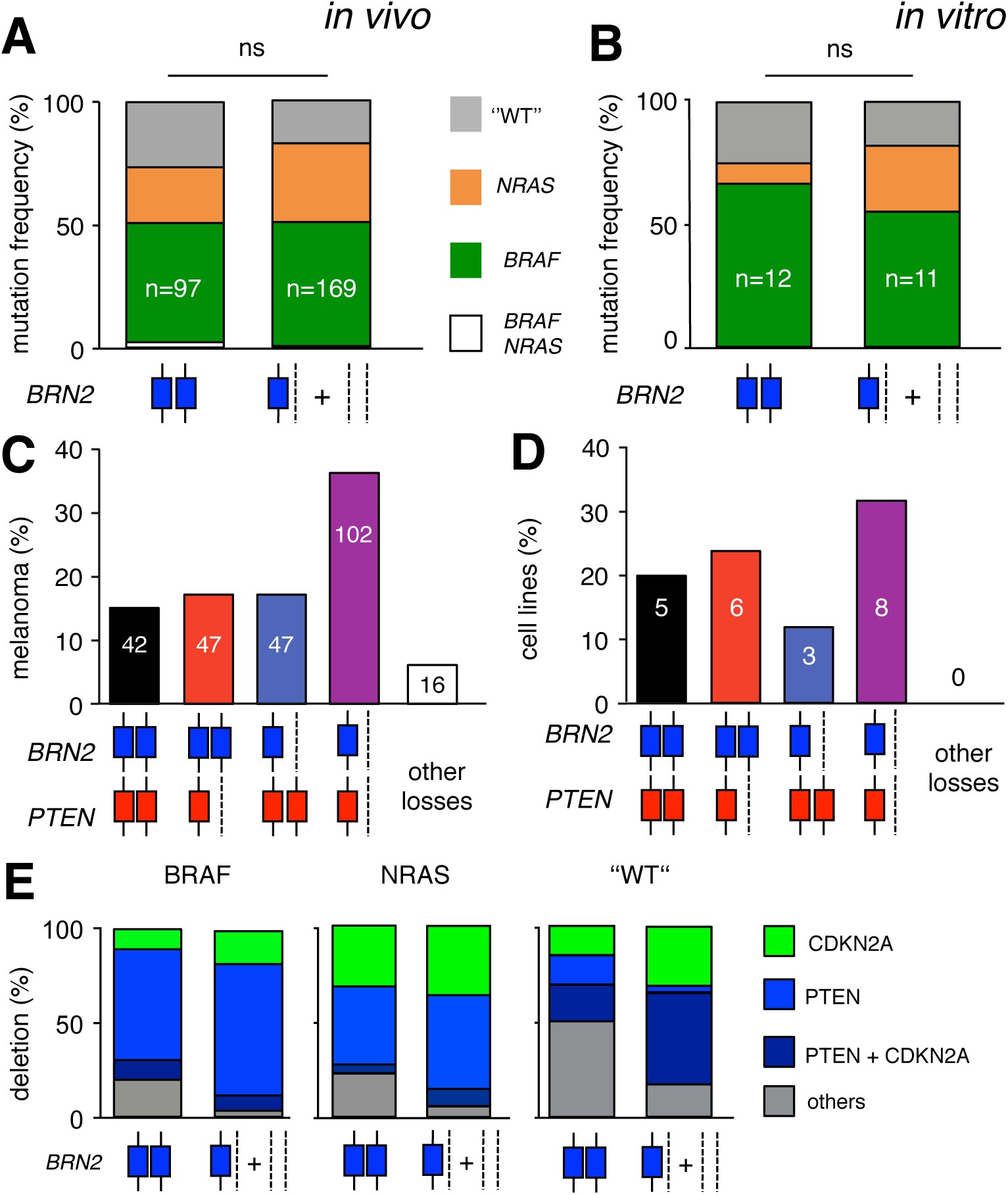
BRN2 is frequently lost in human SKCM and melanoma cell lines independently of NRAS and BRAF driver mutations or CNAs of CDKN2A, and PTEN. (A) Bar graph showing the correlation between the mutation frequency of BRAF (green), NRAS (orange), BRAF + NRAS (white), and “WT” (i.e. non-BRAF/NRAS (grey)) with BRN2 CNAs in human SKCM. CNAs were estimated from GISTIC values. The BRN2 loss group was defined as all samples with mono- or bi-allelic BRN2 loss. BRN2 CNA amplifications (GISTIC > +1) are not shown (n = 21). TCGA sequencing data set (n = 267). Note that from the TCGA_SKCM, only one POU3F2 missense mutation was reported outside of the functional domains. (B) Bar graph showing the correlation of mutation frequency between BRAF (green), NRAS (yellow), or “WT” (grey) with BRN2 CNAs in human melanoma cell lines (n = 23). (C) Bar graph showing the frequency and co-occurrence of BRN2 and PTEN diploidy and mono-allelic loss in human SKCM. CNAs of BRN2 and PTEN are indicated in the pictograms under the graph. TCGA CNA data-set (n = 367. Note there are amplification of PTEN and/or BRN2 in 108 cases. (D) Bar graph showing the frequency and co-occurrence of BRN2 and PTEN diploidy and mono-allelic loss in human melanoma cell lines (n = 23). CNAs of BRN2 and PTEN are indicated in the pictograms under the graph. One cell line showed a mono-allelic gain of PTEN. (E) Bar graph showing the correlation between BRN2 loss and CNAs of CDKN2A, PTEN, and β-catenin in human SKCM samples harboring either BRAF or NRAS mutations or no BRAF/NRAS mutation. CNAs were estimated from GISTIC values. CNAs of BRN2 are indicated in the pictograms under the graph. TCGA CNA data-set (n = 387). BRAF mutants included P318S/L, P367S, G466E, S467L, L485F, G469A/R, N581T, D594N, L597Q, V600K/E/M/G, K601E, E695K, and H725Y. NRAS mutants included G12A, G13D, Q61H/R/K/L, and E62K. PTEN and CDKN2A mutants included full or partial deletions. Statistical analysis was performed using the Chi-square test. ns = non-significant.

**Figure S4.**
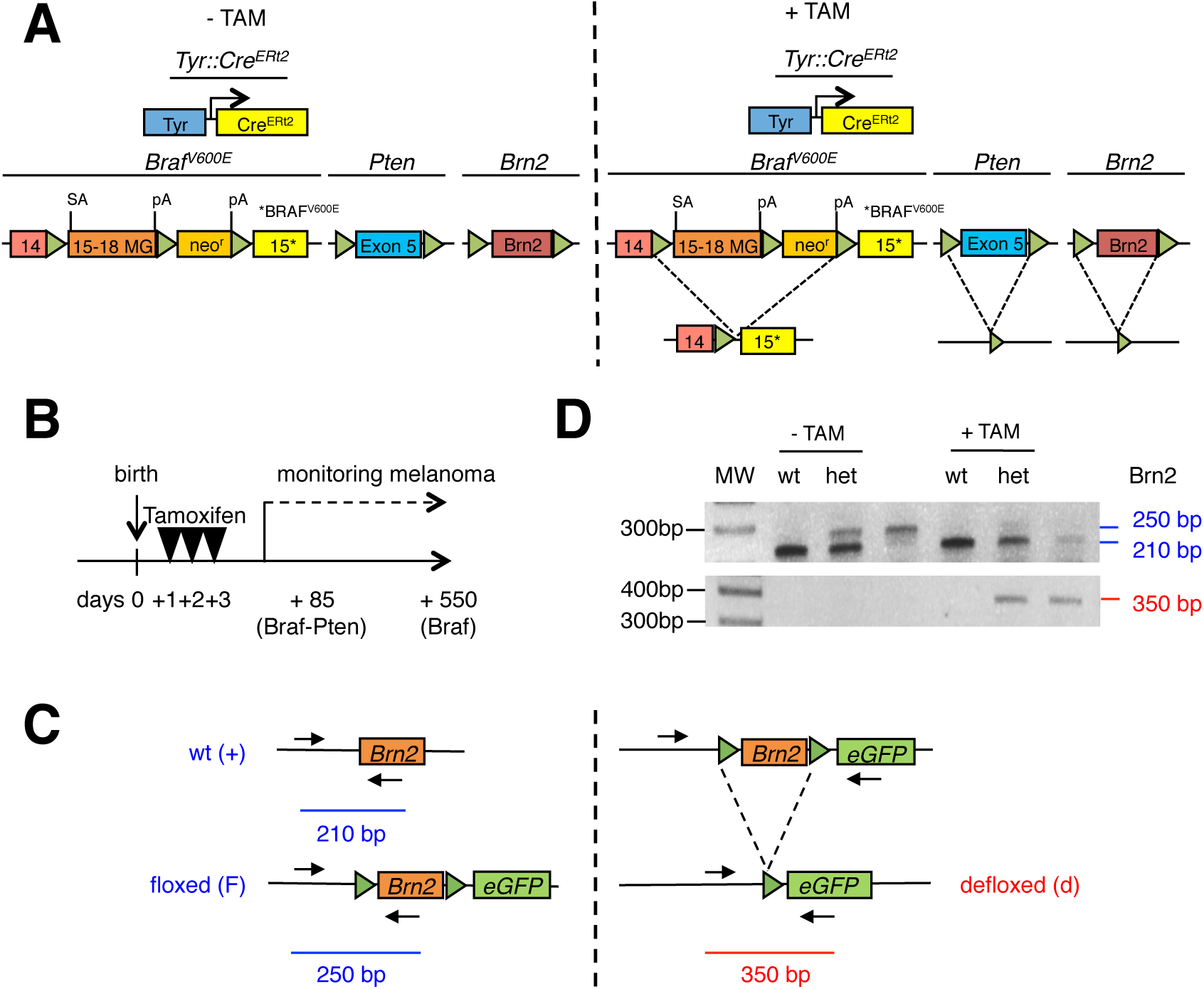
Genetic design and characterization of BRN2 locus knock-out mice. (A) Scheme of the tamoxifen (TAM)-inducible conditional knock-out strategy for Braf, Pten, and Brn2 in the melanocyte lineage. (B) Scheme of TAM application (20 μL/day/mouse [20 μg/mL in DMSO]) for defloxing, leading to the induction of gene knock-out, and mouse follow-up. (C) Scheme of PCR for the detection of wild-type (wt), floxed (f), and defloxed (d) Brn2 alleles in knock-out mice. The arrows indicate the location of the primers and the bars the amplificon length. (D) Representative PCR result for allele identification and defloxing of Brn2. Non-TAM-induced (-TAM) tissue from tails or tissue from TAM-induced tumors (+TAM) from Braf-Pten-WT and Braf-Pten-Brn2 mice.

**Figure S5.**
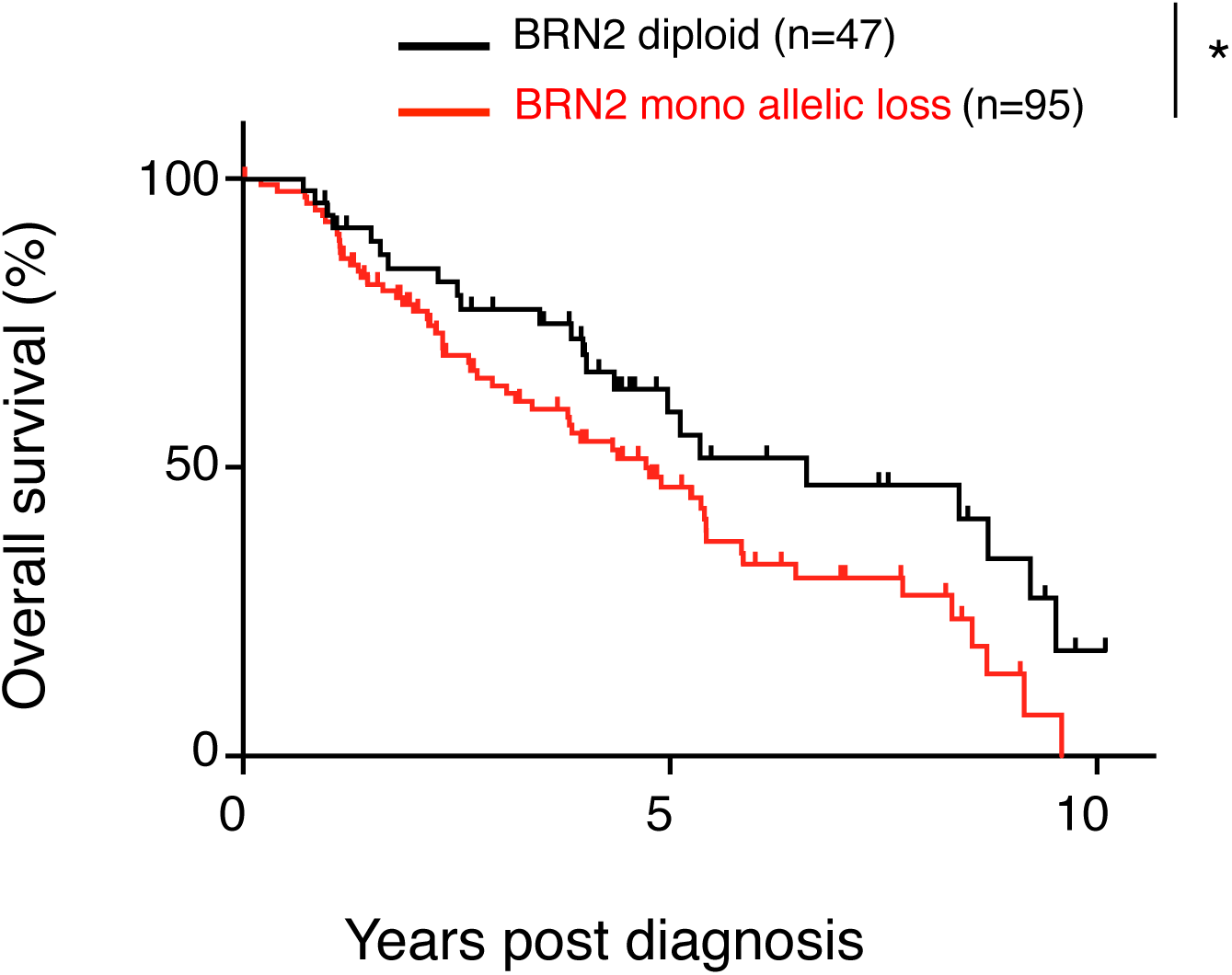
BRN2 loss is associated with reduced overall survival in PTEN mono-allelic patients. Ten-year overall survival of melanoma patients with monoallelic deletion of the PTEN gene according to monoallelic deletion or the absence of deletion of the BRN2 gene. Log-rank (Mantel-Cox) test, *p < 0.05 (p = 0.0352). Two of the patients were bi-allelic.

**Figure S6.**
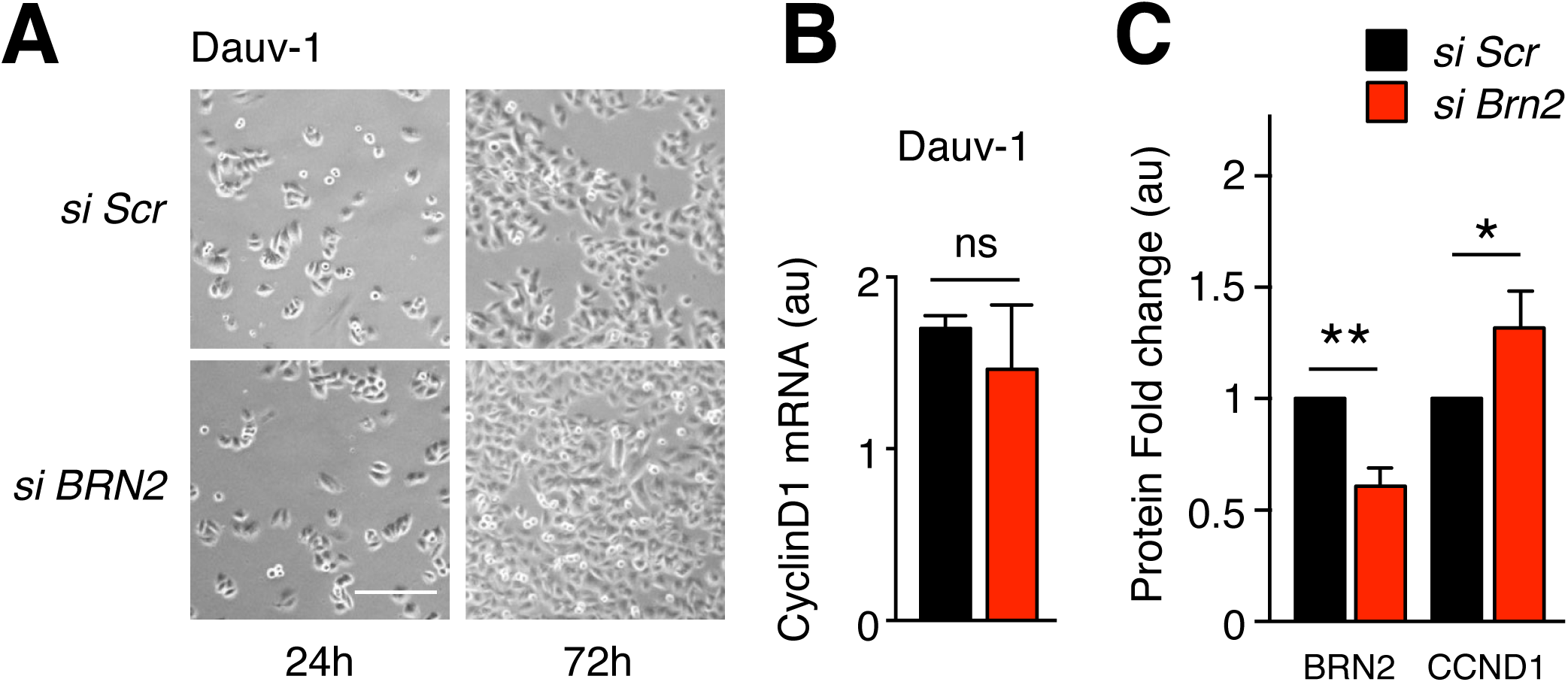
Brn2 loss induces proliferation and increases cyclin D1 protein levels in human melanoma cell lines. (A) Representative photomicrograph of siRNA-mediated knockdown of Brn2 in Dauv-1 human melanoma cell line at 24 and 72 hours after transfection of appropriate siRNA (Scr = scramble). Scale bar = 40 μm. (B) The results of RT-qPCR for cyclin D1 of Dauv-1 human melanoma cells after siRNA-mediated knockdown (n = 3). All values were normalized against those of TBP. a.u.= arbitrary units. (C) Quantification of protein fold change of western blots for Figure 4F. Quantification was performed using Image-J software. All values were normalized against the background and corresponding actin loading control for each sample. Statistical analysis was performed using the unpaired t-test. Error bars correspond to the sem. ns = non-significant, *p < 0.05, **p < 0.01.

**Figure S7.**
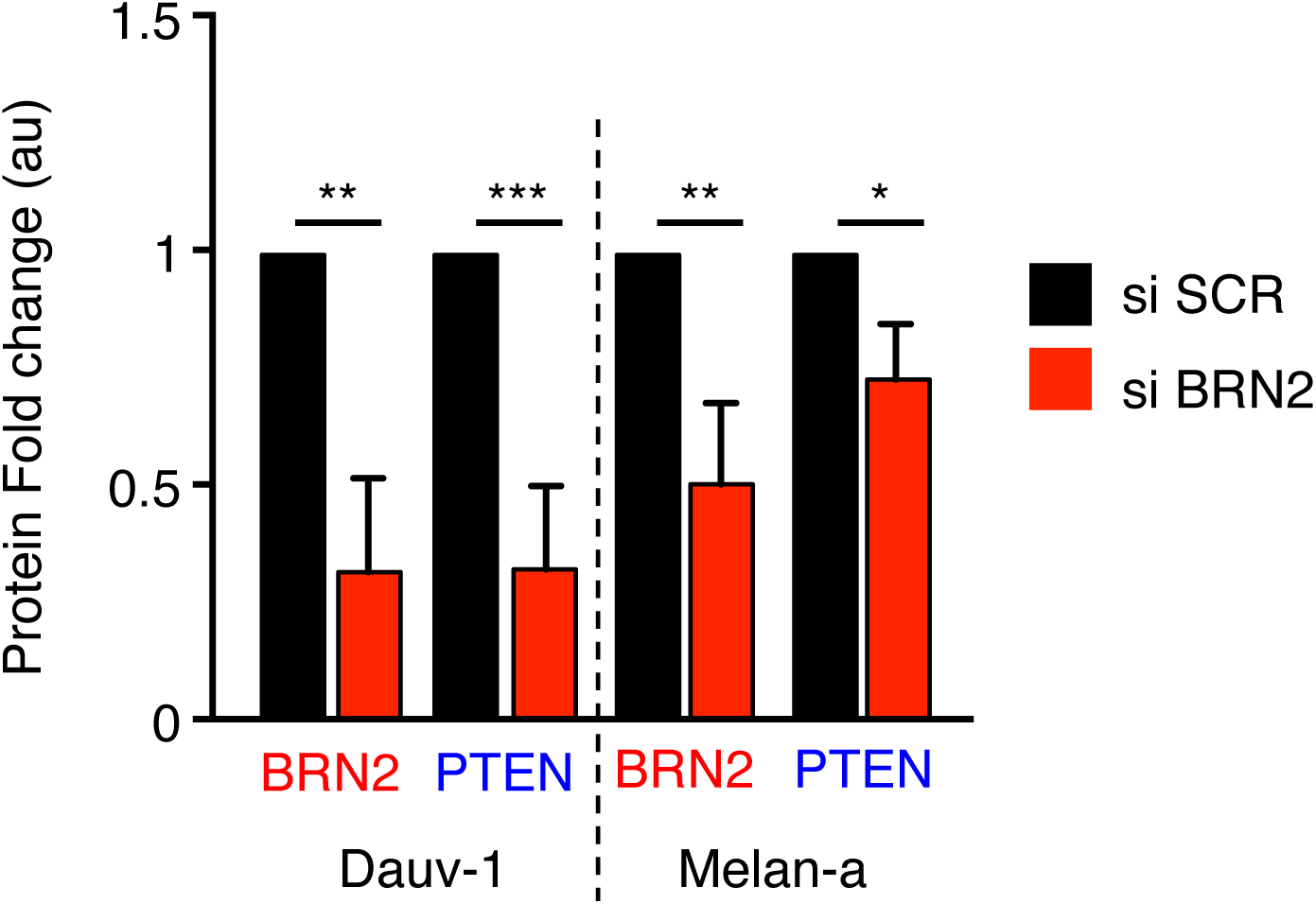
Brn2 knock down reduces Pten level in human melanoma cell lines. Quantification of protein fold change of western blots for Figure 6D. Quantification was performed using Image-J software. All values were normalized against the background and corresponding actin loading control for each sample. Statistical analysis was performed using the unpaired t-test. Error bars correspond to the sem. *p < 0.05, **p < 0.01.

**Figure S8.**
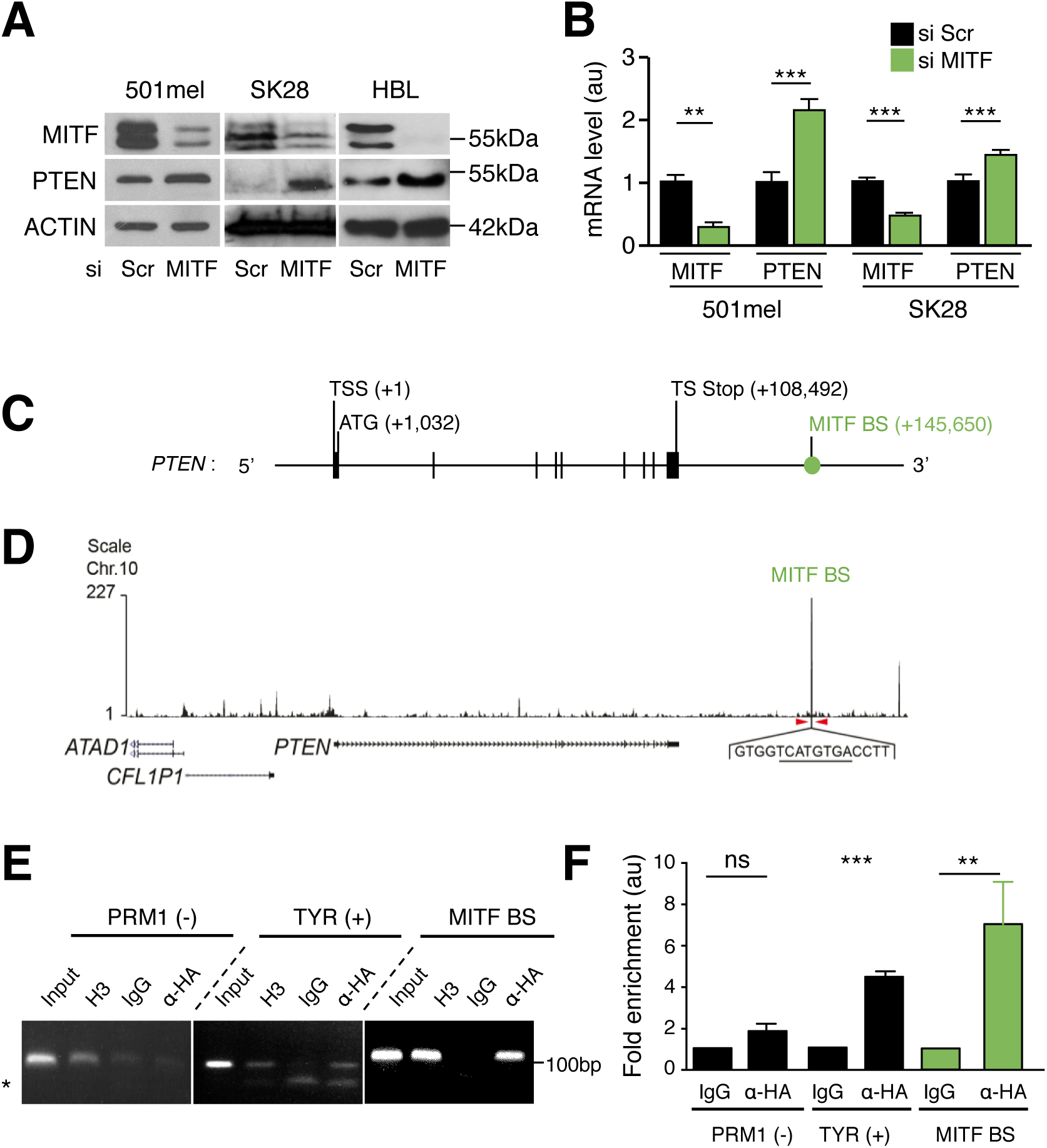
MITF binds downstream of PTEN and MITF loss induces its transcription. (A) Western blot analysis of MITF and PTEN from human melanoma cells (501mel, SK28, and HBL) after siRNA-mediated knockdown. Actin was used as a loading control. A representative western blot is shown. The molecular weight is indicated in kDa. Scr = Scrambled. (B) RT-qPCR of MITF and PTEN from human melanoma cells (501mel and SK28) after siRNA-mediated knockdown. All values were normalized against that of TBP. The analysis was performed on three independent amplifications with technical triplicates, au = arbitrary unit. (C) Scheme of the human PTEN gene containing MITF binding site (BS), represented as green circle. Exons are shown as vertical lines. TSS = transcription start site, and TS Stop = translation stop. All numbering is relative to ATG (+1). (D) Results of genome-wide chromatin immunoprecipitation sequencing (ChIP-Seq) of MITF. PTEN and the adjacent genes are mapped with the arrows indicating strand-orientation and the horizontal rectangles the exons. The peak of MITF binding downstream of human PTEN is indicated by red arrows. The consensus sequence (E/M-Box) is underlined. (E) ChIP assays of MITF binding downstream of PTEN in 501mel human melanoma cells stably expressing HA-Tagged MITF. ChIP assays are performed using an antibody against HA and analyzed after a 30-cycle PCR (exponential phase). The tyrosinase promoter (TYR) and PRM1 were used as positive and negative controls, respectively. MITF BS: MITF binding site downstream of PTEN. Input represents approximately 3% of the input used for the ChIP assay. H3 (histone H3) and IgG (Immunoglobulin G) were used as positive and negative technical controls for each region of interest, respectively. The oligos, their positions on the genome, and sizes of the amplified fragments are shown in Tables S1 and S2. All data shown are representative of three independent assays. *corresponds to the oligonucleotides. (F) Quantification of the ChIP–qPCR is plotted and normalized against IgG as a reference. au = arbitrary unit. PRM1 (-): PRM1 (negative control), TYR (+): tyrosinase promoter (positive control), MITF BS: MITF binding site downstream of PTEN. Statistical analysis was performed using the unpaired t-test. Error bars correspond to the sem. ns = non-significant, *p < 0.05, **p < 0.01, and ***p < 0.001.

